# Local gated-Hebbian learning of deep cerebellar networks with quadratic classification capacity

**DOI:** 10.64898/2026.04.17.718957

**Authors:** Naoki Hiratani

**Affiliations:** Department of Neuroscience, Washington University in St Louis, MO, USA

## Abstract

A central goal of neuroscience is to understand how neural circuit architecture supports learning. While recent work has clarified the computational role of depth in sensory cortical hierarchies, it remains unclear why predominantly feedforward, non-convolutional circuits such as the cerebellum and olfactory system also contain multiple processing layers. Theoretical work in deep learning has shown that two-hidden-layer networks can achieve classification capacity that scales quadratically with the number of intermediate neurons, but these results rely on nonlocal synaptic optimization and are therefore difficult to reconcile with biological learning rules. Here, we show analytically and numerically that a two-hidden-layer network with feedforward gating can achieve quadratic capacity using local three-factor Hebbian learning when intermediate activity is sparse. This architecture supports efficient one-shot learning and, in settings where backpropagation requires many repeated weight updates, offers an advantage in learning speed. Beyond random perceptron tasks, the model also performs well on structured cerebellum-related tasks, including reinforcement-learning-based motor control. Mapping the model onto cerebellar microcircuitry further suggests functional roles for dendritic compartmentalization, branch-specific inhibition, and disinhibitory interneuron pathways. Together, these results extend the Marr-Albus-Ito framework by showing how the presence of multiple intermediate layers in cerebellum-like circuits can support fast, local, and high-capacity learning.

## 1 Introduction

Recent experimental work is rapidly revealing the structure of neural connectivity across both micro- and macroscopic scales [1, 2, 3]. However, we still have limited understanding on functional advantages of particular connectivity motifs or circuit architecture for learning and computation performed by the circuits. Here we address this question focusing on cerebellum, the key regions for motor and cognitive learning [4, 5]. Classical theories often model it as a network with a single intermediate layer [6, 7, 8, 9]. In anatomical terms, however, the cerebellum contains at least two intermediate layers if the cerebellar nuclei are treated as the output layer [4, 10, 11]. Its main feedforward pathway runs from mossy fibers to granule cells, then to Purkinje cells, and finally to the cerebellar nucleus neurons. This makes the canonical cerebellar circuit deeper than is usually assumed in traditional theoretical descriptions. A similar observation applies to insect olfactory systems, which are often compared to the cerebellum because of their related architectural motifs [12, 13, 14]. Especially, in species with multi-glomerular organization, such as locusts, the pathway effectively includes two intermediate layers between sensory neurons and mushroom body output neurons [15]. Nevertheless, the functional role of this depth remains poorly understood. Why should these circuits have multiple intermediate layers at all? What computational advantage, if any, does depth provide in these biological settings?

Recent work in deep learning has offered several possible answers by clarifying when and how depth helps artificial neural networks [16, 17, 18, 19, 20, 21]. One idea is that depth is beneficial when the underlying generative structure of the data is itself hierarchical. For example, in vision, natural scenes have compositional structure, and deep convolutional networks provide an inductive bias that is well matched to that structure [18]. This explanation is compelling in sensory domains such as vision [22], but it is less obviously applicable to olfactory systems, where the generative structure of stimuli is often thought to be relatively shallow. It is also unclear whether this account extends to the cerebellum.

A second possibility is that depth helps by introducing an implicit simplicity bias during learning, even in otherwise standard feedforward networks without convolutional structure [16, 23]. In this view, deeper networks can be easier to train toward useful solutions because gradient-based learning favors certain classes of functions. However, this explanation depends on learning dynamics driven by backpropagation algorithm (below we refer it as ‘backprop’ to avoid confusion with dendritic backpropagation), which is considered biologically implausible [24]. For that reason, it is also unlikely to explain why multilayer architectures are found in cerebellar-like circuits.

A third hypothesis is that depth increases memory capacity. It has long been conjectured [25], and more recently proven [19, 20, 21], that in networks with two intermediate layers, perceptron capacity, the number of random patterns the network can correctly classify, can scale as *L*_1_*L*_2_ where *L*_1_ and *L*_2_ are the numbers of neurons in the first and second intermediate layers, respectively. This scaling is dramatically larger than in the classical Marr-Albus-Ito framework, where capacity scales only with the number of granule cells. This idea is especially relevant for the cerebellum and olfactory systems, both of which are thought to learn mappings from high-dimensional inputs to behaviorally meaningful outputs.

At the same time, existing mechanisms proposed for achieving this quadratic capacity depend on highly specialized constructions of the weight matrices [19, 21]. Such constructions are difficult to reconcile with biological constraints and are unlikely to be implemented literally in the brain. As a result, it remains unclear whether having two intermediate layers is sufficient to support quadratic capacity under biologically realistic conditions, and whether this could help explain the organization of cerebellar circuitry.

In this work, we first show both analytically and numerically, that by adding gating into the circuit, the biologically-plausible three-factor Hebbian learning achieves the quadratic capacity. Moreover, due to the Hebbian update, this model achieves one-shot learning of associations unlike the traditional backprop-based or perceptron-like learning where multiple presentation of a single input-output pair is necessary for learning of correct classification. We also show that this model achieves fast, high-capacity learning of motor control tasks in continual action space, supporting its applicability to fine spatio-temporal mapping relevant to cerebellum function, beyond simple classification problem.

To establish the functional advantage of this circuitry, we map the model onto the cerebellar circuit. Although the naive model does not fully match cerebellar anatomy, a version constrained by known cerebellar connectivity still supports high-capacity learning in a pattern-classification task. This mapping further suggests functional roles for Purkinje cell dendritic computation and disinhibitory connectivity motifs within the cerebellar microzone. Moreover, the model makes a testable prediction about the fine-scale dendritic connectivity between granule cells and Purkinje cells. Parts of this work were previously reported in abstract form [26].

## 2 Results

### 2.1 Classification capacity of networks with zero, one, and two intermediate layers

We begin by briefly recapping the perceptron classification problem and its recent developments. In the perceptron setting, the goal is to classify neural activity patterns ***x***_1_, ***x***_2_, …, ***x***_*P*_ according to their corresponding targets *y*_1_, *y*_2_, …, *y*_*P*_ (Fig. 1A). For example, in the cerebellum, ***x*** and *y* may correspond to cortical representations of sensory inputs and the corresponding motor reflex outputs, respectively. In olfactory systems, ***x*** corresponds to the representation of olfactory inputs at the sensory periphery, whereas *y* represents odor valence.

**Figure 1:**
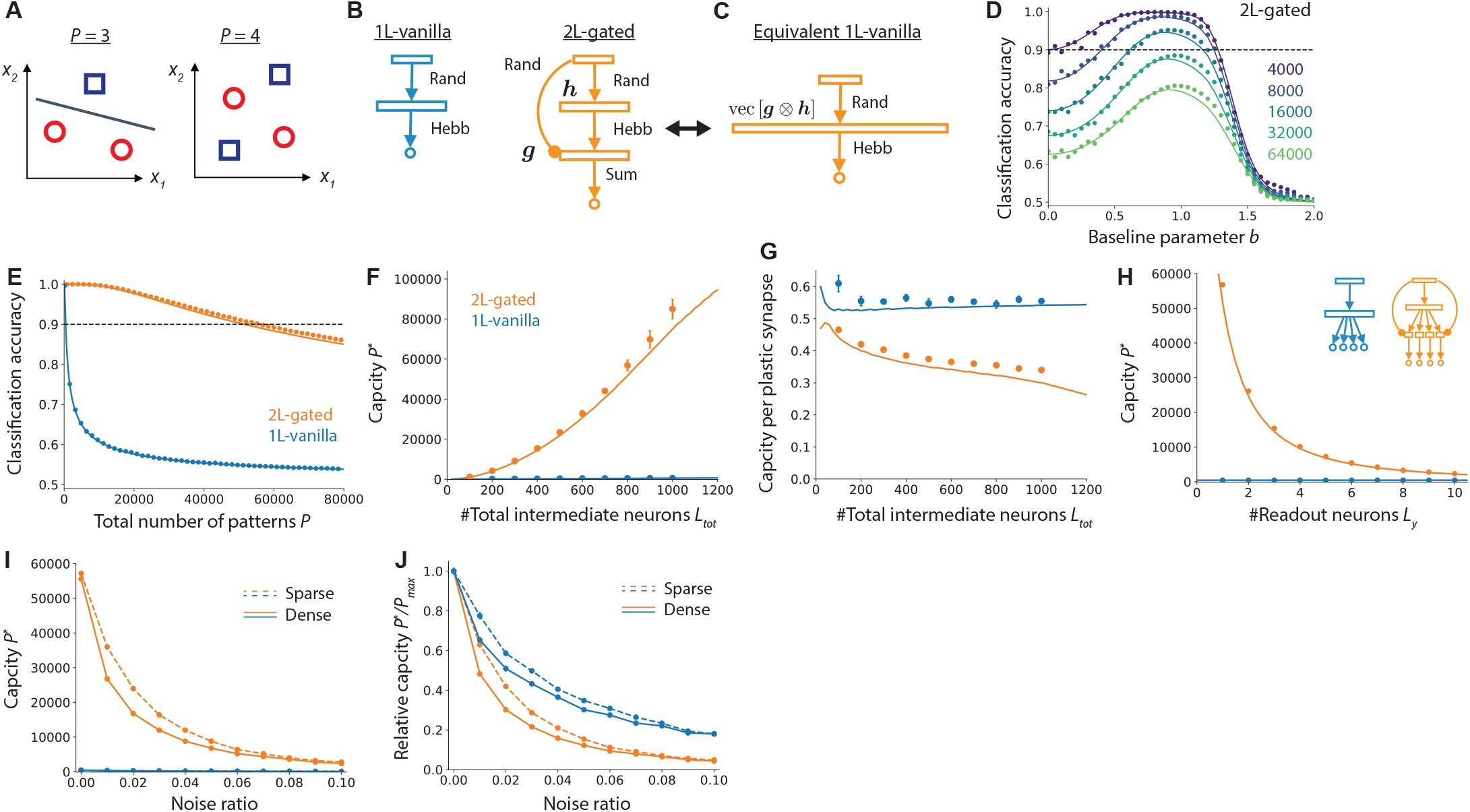
**A)** Schematic of the perceptron problem. **B)** Illustration of the one-layer vanilla model (1L-vanilla; left) and the two-layer gated network model (2L-gated; right). Rand: random weights; Hebb: Hebbian weights; Sum: summation (i.e., constant positive weights). **C)** The equivalent one-layer network of a two-layer gated network. vec[***g*** ⊗ ***h***] represents the flattened vector of the outer product of two vectors ***g*** and ***h*. D)** Classification accuracy under different values of the baseline parameter *b* in the two-layer gated model with *L*_0_ = 100 and *L*_1_ = *L*_2_ = 400. Large *b* makes activity in the intermediate layers sparse, whereas *b* = 0 means that, on average, half of the neurons are active. **E)** Classification accuracy of the two models under the optimal baseline parameter *b*^∗^ and *L*_*tot*_ = 800 (i.e., *L*_1_ = 800 for 1L-vanilla; *L*_1_ = *L*_2_ = 400 for 2L-gated). The horizontal dashed line indicates the threshold used to estimate capacity. **F)** Capacity as a function of the total number of neurons in the intermediate layers, *L*_*tot*_. Error bars represent the standard error of the mean estimated with bootstrap over 10 random seeds, unless otherwise stated. **G)** Same as panel F, but with capacity normalized by the number of plastic synapses. **H)** Capacity in the presence of multiple readout neurons under *L*_*tot*_ = 800. **I**,**J)** Capacity under *L*_*tot*_ = 800 in the presence of input noise. Solid lines represent the full model, whereas dashed lines show models in which the random connections (Rand in panel B) are sparsified with connection probability *ρ*_*c*_ = 0.3. In panels C–G, lines indicate theoretical results and points indicate numerical results. In panels I–J, lines are linear interpolations. See Methods Sections 4.2.1 and 4.2.2 for model details.

If we consider linear classification of input activity patterns, *y* = *σ*(***w*** · ***x***), the number of patterns that can be classified scales with *L*_0_, where *L*_0_ is the number of neurons in the input layer [27]. For example, when there are only two input neurons (*x*_1_ and *x*_2_ in Fig. 1A), up to three patterns can be classified, but when there are four patterns, linear classification is no longer guaranteed [28].

If we add one intermediate layer by randomly projecting the input, the classification capacity instead depends on the size of the intermediate layer, because the final linear readout is now performed from that layer [29, 30]. Denoting the number of neurons in the intermediate layer by *L*_1_, the capacity scales with *L*_1_. Thus, if the intermediate layer contains many more neurons than the input layer (i.e., *L*_1_ ≫ *L*_0_), classification capacity can in principle be increased by a factor of *L*_1_*/L*_0_. This idea has been proposed as an explanation for the large number of neurons in the cerebellar granule cell layer [6, 7] and in the Kenyon cell layer of insect olfactory circuits [31, 32, 33]. However, this improvement is only linear in the number of intermediate-layer neurons, implying that a very large number of neurons is required to achieve high capacity.

By contrast, it has recently shown that, in networks with two intermediate layers, classification capacity can scale with the product of the two layer sizes, *L*_1_*L*_2_, where *L*_1_ and *L*_2_ are the numbers of neurons in the first and second intermediate layers, respectively [19, 20, 21]. This implies that, if the total number of intermediate neurons is constrained to *L*_*tot*_ by energetic and spatial costs [34], splitting them into two layers yields a capacity scaling of 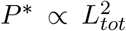. If the same neurons are instead placed in a single layer, the capacity scales only as 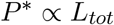. Thus, two intermediate layers can in principle increase classification capacity quadratically.

As discussed above, however, previously proposed constructions of two-hidden-layer networks with quadratic capacity rely on elaborate schemes that are difficult to implement in a biologically plausible manner, leaving their biological relevance unclear. Although such networks might be implemented using backpropagation or more biologically plausible surrogates, backpropagation-based learning is typically slow and requires repeated presentations of the same stimulus, and biologically plausible surrogates often face additional limitations in learning speed and performance [35, 24]. This contrasts with the rapid, often one-shot learning observed in the brain. Slow learning is especially problematic in high-capacity systems, because high capacity is of limited use if the network takes too long to approach it. It therefore remains unclear whether rapid local learning can coexist with high-capacity classification.

### 2.2 Gated two-intermediate-layer networks achieve quadratic capacity with three-factor Hebbian learning

Can quadratic capacity be achieved in a biologically plausible manner? And if so, can it be achieved with one-shot learning? Here we show, both analytically and numerically, that a network with two intermediate layers and feedforward gating supports one-shot Hebbian learning for pattern classification and achieves capacity that scales quadratically with the number of neurons in the intermediate layers.

The orange model in Fig. 1B (right) illustrates the proposed architecture. The activities of the intermediate-layer units *h*_*i*_ and *g*_*i*_, the second intermediate-layer units *z*_*i*_, and the output unit *y* follow (see Methods Section 4.2.1 for details):

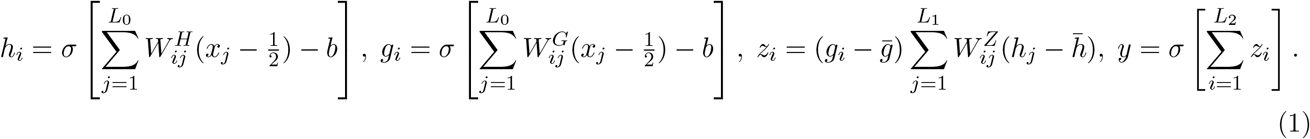

In the cerebellar interpretation of this model, the four layers correspond to mossy fibers (*x*_*i*_), granule cells (*h*_*i*_), Purkinje cells (*z*_*i*_), and a cerebellar nucleus neuron (*y*; see Fig. 5 for the detailed mapping). The first intermediate layer *h*_*i*_ is analogous to the intermediate layer in the traditional Marr–Albus model. The matrix 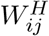 randomly expands the input ***x*** = [*x*_1_, …, *x*_*N*_], and the nonlinear function *σ*, defined here as the Heaviside function (*σ*(*x*) = 1 if *x >* 0 and 0 otherwise), ensures a nonlinear transformation of the input. The gating unit *g*_*i*_ multiplicatively modulates the activity of the second intermediate layer *z*_*i*_, which could be implemented through shunting inhibition. Parameter *b* is the baseline controlling the sparsity of intermediate layer activity, and 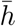 and 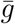 are average activity at units *h* and *g*. We set *W*^*H*^ and *W*^*G*^ to be random, whereas *W*^*Z*^ is learned according to

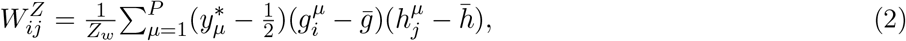

where *Z*_*w*_ is a normalization constant, and 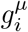 and 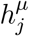 denote the activities of units *g*_*i*_ and *h*_*j*_ for pattern ***x*** = ***x***_*µ*_. We refer to this weight update following the association between the target 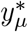 and activities 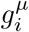 and 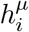, as Hebbian, to disambiguate it from the error-based learning rule introduced later (Eq. 5).

In this setting, for random input patterns, the output noise-to-signal ratio approximately follows (see Methods Section 4.3)

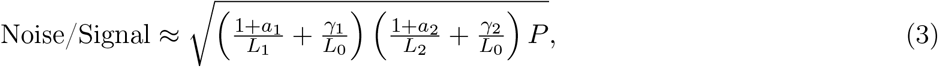

where (*a*_1_, *γ*_1_) and (*a*_2_, *γ*_2_) are factors that depend on the sparsity of *h*_*i*_ and *g*_*i*_, respectively, and recall that *L*_1_ and *L*_2_ are the number of neurons in the first and the second intermediate layers. In the sparse limit, where *a*_1_, *γ*_1_, *a*_2_, and *γ*_2_ all approach zero (see Fig. 6 in Methods Section 4.3.4), the noise-to-signal ratio becomes 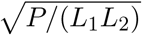, indicating that the number of patterns that can be classified scales as *P* ∝ *L*_1_*L*_2_. Hence, the classification capacity of this network scales quadratically with the number of neurons in the intermediate layers.

Intuitively, this capacity expansion arises because multiplicative gating effectively creates a one-layer network with an intermediate representation of size *L*_1_*L*_2_. Specifically, from Eq. 1, the output can be rewritten as

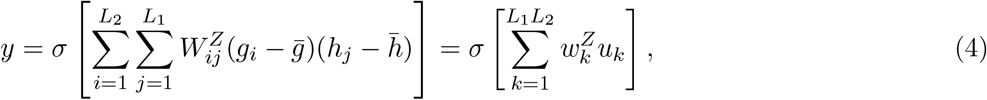

where 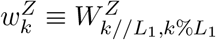 and 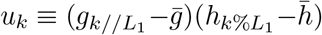 (Fig. 1C; in vector form: 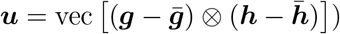.

Because the capacity of a one-intermediate-layer network scales with the size of its intermediate representation, this construction enables *P* ∝ *L*_1_*L*_2_ capacity when *g* and *h* are sparse and thus almost uncorrelated.

We numerically simulated this network using randomly generated patterns and refer to it as the 2L-gated model. For comparison, we also considered the classical Marr–Albus-type one-intermediate-layer network, in which the first set of connections is random and the second set is learned with a Hebbian-type association between the target and the input. We refer to this model as the 1L-vanilla model (see Methods Section 4.2.2). Additional variants are considered in Fig. 2.

**Figure 2:**
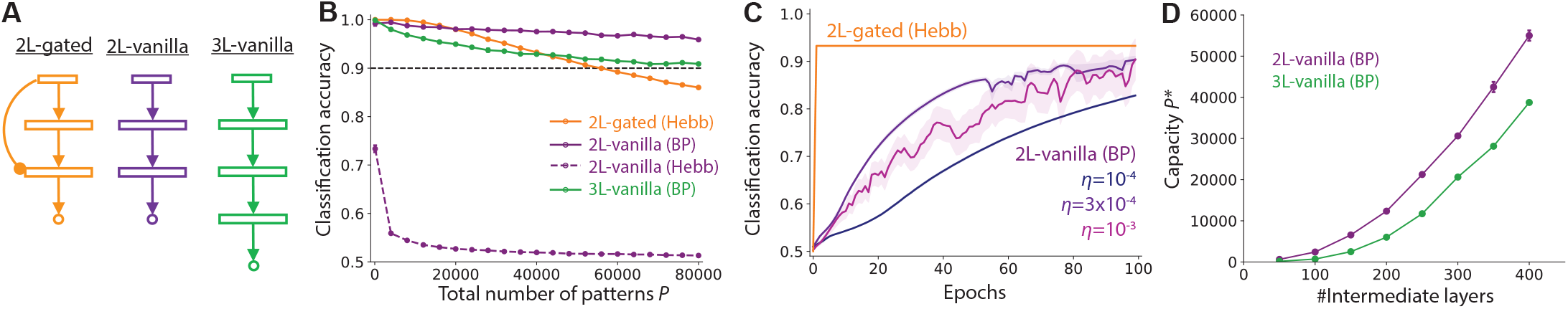
**A)** Illustration of the proposed two-layer gated model (left) and two alternative models: a two-layer vanilla model (middle) and a three-layer vanilla model (right). **B)** Classification accuracy of the 2L-gated model with Hebbian learning, the 2L-vanilla model with BP (backprop) training (purple solid), the 2L-vanilla model with Hebbian learning (purple dashed), and the 3L-vanilla model with BP training (green). **C)** Comparison of learning speed between Hebbian learning in the 2L-gated model and BP training of the 2L-vanilla model under *P* = 40000. The three curves for the 2L-vanilla model represent learning curves under different learning rates. In panels B and C, we set the total number of neurons in the intermediate layers to *L*_*tot*_ = 800 by setting *L*_1_ = *L*_2_ = 400 for the two-layer models and *L*_1_ = *L*_2_ = *L*_3_ = 266 for the three-layer model. We implemented backprop using the Adam optimizer [38] with default hyperparameters (*β*_1_ = 0.9, *β*_2_ = 0.999) with mini-batch where mini-batch size was set to *P/*100. Error bars are the standard error of the mean over 10 random seeds. **D)** Capacity of two- and three-layer networks trained with backpropagation as a function of the total number of neurons in the intermediate layers, *L*_*tot*_. See Methods Section 4.2.3 for details of the 2L/3L-vanilla models.

As previously observed in one-intermediate-layer networks [6, 30], classification accuracy in the 2L-gated network depends strongly on the baseline parameter *b*, which controls the sparsity of intermediate-layer activity (Fig. 1D). We therefore optimized this bias when estimating classification capacity below. Under the optimal baseline parameter *b*, classification accuracy in the 2L-gated model remained high even when the network stored ~ 60,000 patterns (orange line in Fig. 1E; points: numerical simulations; line: theory from Section 4.3.2; we set the input layer width to *L*_0_ = 100 and the intermediate layer width to *L*_1_ = *L*2 = 400). By contrast, the 1L-vanilla model showed a rapid decay in classification accuracy as the number of stored patterns increased (blue line; *L*_1_ = 800).

As predicted, the capacity of the 2L-gated model increased quadratically with the total number of hidden neurons (orange points in Fig. 1F). Here, we defined capacity as the largest number of patterns for which the network achieved 90% classification accuracy (black dashed line in Figs. 1D–E). By contrast, the 1L-vanilla model, corresponding to the Marr–Albus framework, achieved a much smaller capacity (blue points in Fig. 1F). For this comparison, we fixed the total number of intermediate-layer neurons across models. Thus, under *L*_*tot*_ = 800, the 1L-vanilla model contains *L*_1_ = 800 neurons in its intermediate layer, whereas the 2L-gated model contains *L*_1_ = *L*_2_ = 400 in its two intermediate layers. The contrast between the blue and orange curves in Fig. 1F illustrates the functional advantage of the proposed framework over the traditional Marr–Albus-type cerebellar model.

The key to this dramatic improvement is that learning occurs between the two intermediate layers, yielding 𝒪(*L*_1_*L*_2_) plastic synapses. By contrast, in the 1L-vanilla model, only 𝒪(*L*_1_) output synapses are available for learning, resulting in a much lower capacity. Indeed, when capacity is normalized by the number of plastic synapses, the 1L-vanilla model robustly achieves higher capacity per synapse (blue vs. orange curves in Fig. 1G). Moreover, when there are multiple readouts each performs independent classification, the 1L-vanilla model can readily scale without sacrificing capacity (blue model and line in Fig. 1H; the blue line is horizontally flat). By contrast, the capacity of the 2L-gated model decreases reciprocally with the number of independent readouts (orange line), because each output target requires a dedicated hidden-layer module. Nevertheless, because its original capacity is much larger, the 2L-gated network still retains an advantage over the 1L-vanilla model even with ten readouts. Notably, anatomical evidence indicates that Purkinje cells within the same cerebellar microzone tend to project to the same set of cerebellar nucleus neurons [36, 37], implying that the effective output dimensionality per microzone is likely to be small.

The proposed model is less robust to input noise than one-hidden-layer models (orange vs. green solid lines in Fig. 1I,J; panel J shows capacity relative to the noiseless condition). Nevertheless, simply sparsifying the random connections *W*^*H*^ and *W*^*G*^ substantially improved robustness (dashed vs. solid lines in Fig. 1I; the connection probability was set to *ρ*_*c*_ = 0.3 and *ρ*_*c*_ = 1.0 for dashed and solid lines, respectively), as suggested previously [13].

### 2.3 Gating enables high-capacity Hebbian learning

The results above show that, in a gated two-layer network, biologically plausible three-factor Hebbian learning can achieve quadratic capacity. This raises two related questions: whether gating is necessary for achieving quadratic capacity, and whether there is any benefit to making the network even deeper.

Regarding the first question, when trained with backprop, a two-layer network without gating was sufficient to achieve a capacity higher than that of the gated network trained with the Hebbian rule (solid purple vs. orange in Fig. 2B; see Methods Section 4.2.3 for details). However, when the number of patterns was large, backprop-based training of the two-layer network required many weight updates over repeated presentations of the same patterns (blue, purple, pink lines in Fig. 2C; the three curves represent learning trajectories under different learning rates of the Adam optimizer [38] we used for backprop). By contrast, the proposed two-layer gated network achieved high capacity after only a single training epoch, due to the one-shot nature of the Hebbian update (orange line). This suggests a potential advantage of the gated network over the non-gated network in terms of learning efficiency.

When Hebbian-type associative learning between the input and the target was applied to the 2L-vanilla network, the capacity was substantially lower than that achieved with backpropagation training (dashed versus solid purple lines). This reduction occurs because, without gating, a naive local learning rule causes neurons in the second intermediate layer to develop similar weight structures. This redundancy in the learned weights limits classification capacity, although it may help denoise activity in the presence of neural noise [11]. Thus, without gating, the network either requires slower and less biologically plausible learning or exhibits a marked reduction in capacity, supporting the functional importance of gating.

The second question concerns the possible benefit of greater depth in cerebellum-like networks. When we added one more layer while keeping the total number of neurons in the intermediate layers fixed, the resulting three-layer model performed worse than the two-layer model under backpropagation training (green vs. purple in Figs. 2B,D). This can be understood from the fact that capacity scales approximately with the total number of plastic synapses [21, 25]. For a network with *k* intermediate layers, the total number of synapses between intermediate layers is 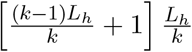, which is maximized at *k* = 2.

### 2.4 Application to structured tasks

The results above demonstrate an advantage of the 2L-gated model for classifying random input-output pairs, but this analytically tractable setting remains idealized and does not fully capture the kinds of learning problems faced by the cerebellum. While the cerebellum takes part in diverse learning tasks, those tasks likely involve structured, non-random inputs; error-driven learning rather than purely associative learning; spatiotemporal task structure; and motor control [5, 14, 40, 41]. To examine whether the proposed model remains effective in such settings, we applied it to two additional classes of problems below.

First, to test its applicability to error-driven learning from structured inputs, we applied the model to Omniglot, a handwritten character classification dataset [39]. We randomly assigned binary labels (0 or 1) to 1024 characters selected from 50 writing systems, and then trained the network to predict the assigned label from the image. We evaluated the trained network using both memorization and generalization tests (Fig. 3A), corresponding to accuracy on the training images and on held-out images of the same characters, respectively (see Methods Section 4.1.2 for details). To mimic cortical preprocessing of sensory inputs before projection to the cerebellum, we first passed the images through a pre-trained convolutional network trained with self-supervised learning [42, 43], and then projected its output to the proposed two-layer gated model and to the control models (Fig. 3B).

**Figure 3:**
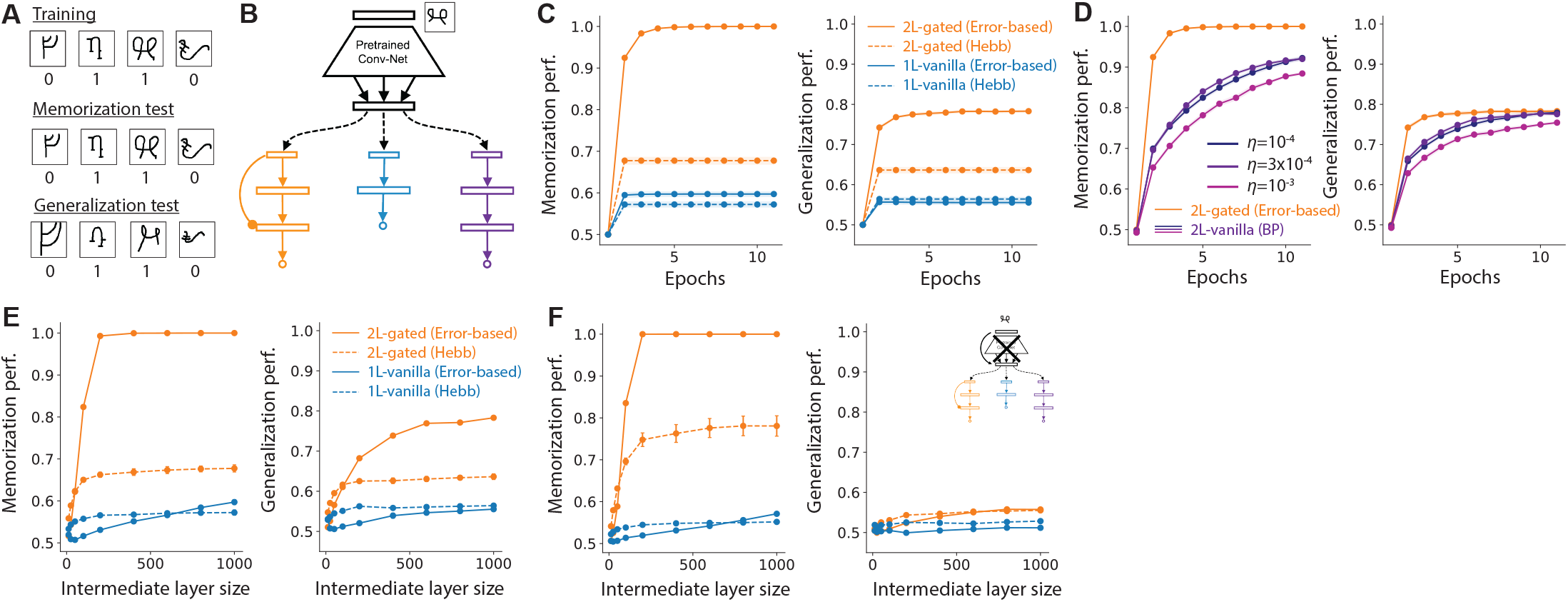
**A)** Schematic of the task. We randomly assigned binary labels to 1024 characters from the Omniglot dataset [39], and then trained the network to associate images with their corresponding labels. We evaluated memorization and generalization using images from the training set and held-out images of the same characters, respectively. **B)** We first processed the images using a convolutional network pre-trained with a self-supervised learning algorithm (corresponding to cortex), and then passed the resulting representation to the proposed two-layer gated model (2L-gated; orange), the one-layer Marr–Albus-type model (1L-vanilla; blue), and the two-layer vanilla model (2L-vanilla; purple). See Methods Section 4.1.2 for task details and pre-training procedures. **C)** Memorization and generalization accuracy during repeated training on the same dataset for the 2L-gated and 1L-vanilla models under *L*_*tot*_ = 2000. Solid and dashed lines represent error-based and Hebbian-type learning, respectively (Eqs. 5 and 2). The x-axis shows the number of training epochs. By construction, performance under the Hebbian update saturates after one epoch. **D)** Comparison of learning curves between the 2L-gated model trained in an error-driven manner and the 2L-vanilla model trained with backprop under various learning rates. **E)** Model performance after 10 epochs of training as a function of the total number of neurons in the intermediate layers, shown for the 2L-gated and 1L-vanilla models. **F)** Model performance in the absence of the pre-trained convolutional network. As expected, the model fails to generalize while retaining memorization performance.

To capture the error-driven nature of learning, we modified Eq. 2 as follows:

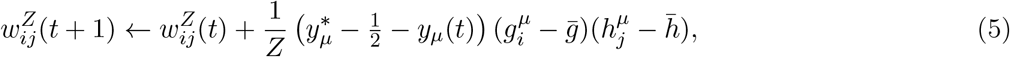

where the *y*-dependent term includes subtraction of the current prediction *y*_*µ*_(*t*), making the update error-driven rather than purely associative. We found that this error-based iterative formulation improved performance on this task in terms of both memorization and generalization (solid orange vs. dashed orange lines in Fig. 3C). Moreover, as in the perceptron setting, the 2L-gated network outperformed the 1L-vanilla network with the same number of intermediate-layer neurons in both memorization and generalization (orange vs. blue lines in Fig. 3C,E). These results suggest that the advantage observed in the analytically tractable perceptron setting (Fig. 1) extends to error-driven learning from structured image inputs.

Furthermore, the 2L-gated network learned faster than the two-layer vanilla network trained with backprop (Fig. 3D; the three purple curves correspond to different learning rates). Although the 2L-gated network achieved higher memorization performance (Fig. 3D, left), this did not come at the expense of generalization relative to the two-layer vanilla network (Fig. 3D, right). Notably, generalization performance dropped sharply when the pre-trained network was removed and the raw input was projected directly to the cerebellum-like network (Fig. 3F vs. E), suggesting the benefit of preprocessed cortical inputs to cerebellum-like networks when generalization is required.

We next considered reinforcement learning (RL) benchmarks to examine whether the model can also handle tasks involving spatiotemporal structure and sensorimotor control, in addition to structured inputs and error-driven learning.

To this end, we studied two simple RL tasks, Reacher and Pendulum (`reacher_v5` and `pendulum_v1` in Gymnasium [44]). We implemented these tasks using an actor–critic framework in which both the actor and critic were feedforward neural networks [45, 46, 47]. In this mapping, the actor corresponds to the cerebellum, whereas the critic corresponds to the basal ganglia. We trained the models using Proximal Policy Optimization (PPO) [45], a standard deep RL algorithm for continuous action spaces, except that, rather than training the full network end-to-end with backprop, we updated the 2L-gated and 1L-vanilla actors using local synaptic update rules driven by the error signal generated by PPO (see Methods Section 4.4.4). We trained the critic with backprop, because our main question here was whether a cerebellum modeled as a 2L-gated network can support motor learning, and if so, whether it performs better than the Marr–Albus-type 1L-vanilla model.

We first applied these models to the Reacher task, in which the goal is to control the torques of a two-joint robot arm so that it reaches a target. This is a relatively simple task because feedback is dense. Under this condition, models with 2L-gated and 1L-vanilla actors both performed comparably to the 2L-vanilla network trained with backprop (Fig. 4B). We then applied the same models to the Pendulum task, in which the goal is to swing the pendulum upright while using minimal torque. In this setting, both the 2L-vanilla and 2L-gated networks quickly learned to perform the task. By contrast, the 1L-vanilla network, despite having the same total number of intermediate-layer neurons, struggled to master the task (cyan vs. orange and purple in Fig. 4C). These results suggest that the two-layer gated network is applicable to spatiotemporal control tasks and can outperform the Marr–Albus-type network in at least some settings. Overall, these results indicate that the potential advantage of the proposed model is not limited to the random perceptron setting.

**Figure 4:**
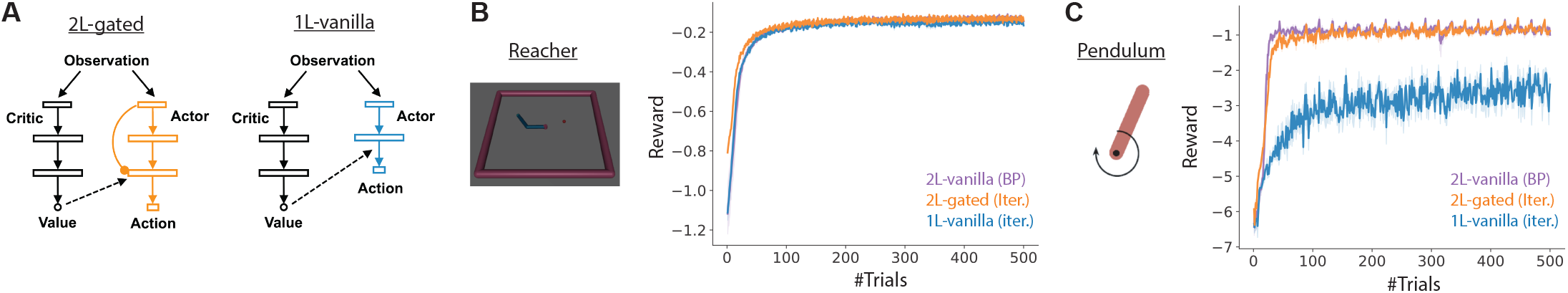
**A)** Schematic of the actor–critic deep RL models. We modeled the actor network as either a two-layer gated network (2L-gated), a one-layer vanilla network (1L-vanilla), or a two-layer vanilla network (2L-vanilla; control). For the critic network, we used the two-layer vanilla network in all settings. **B)** Learning curves of the 2L-gated (orange), 1L-vanilla (cyan), and 2L-vanilla (purple) networks in the Reacher task (illustrated on the left). The goal of the task is to touch the target by controlling the torque applied to a two-joint robot arm. The purple line corresponding to the 2L-vanilla model is hidden under the cyan line. **C)** Learning curves of the three networks in the Pendulum task, where the goal is to swing the pendulum to the upright position using minimal torque. In both panels B and C, the y-axis shows the average reward per trial, averaged over 5 random seeds, and the shaded areas represent the standard error of the mean. We optimized the learning rate of each neural architecture to maximize cumulative reward, while keeping the other hyperparameters fixed. See Methods Section 4.4.4 for details.

### 2.5 Mapping the two-layer gated model to cerebellar circuitry

The results above suggest that the proposed two-layer gated model is biologically plausible and exhibits strong learning performance. To examine whether it provides a functional model of cerebellar circuitry, and to gain insight into cerebellar organization, we next map this model onto the mammalian cerebellum.

As illustrated in Fig. 5A (left), the four layers of the two-layer gated model map naturally onto mossy fibers (MF), granule cells (GC), Purkinje cells (PKC), and cerebellar nucleus (CN) neurons, respectively, thereby capturing the basic feedforward organization of the cerebellum. In particular, unlike traditional one-intermediate-layer models, this framework explicitly incorporates the cerebellar nuclei as the output unit of the cerebellum. The convergence of Purkinje cells onto cerebellar nucleus neurons is broadly consistent with cerebellar microzone organization, in which nearby Purkinje cells often converge onto the same cerebellar nuclear module [36, 37]. Moreover, in the model, plasticity occurs at the GC–PKC connections under modulation by supervised signals, consistent with inferior-olive-driven plasticity at GC–PKC synapses.

**Figure 5:**
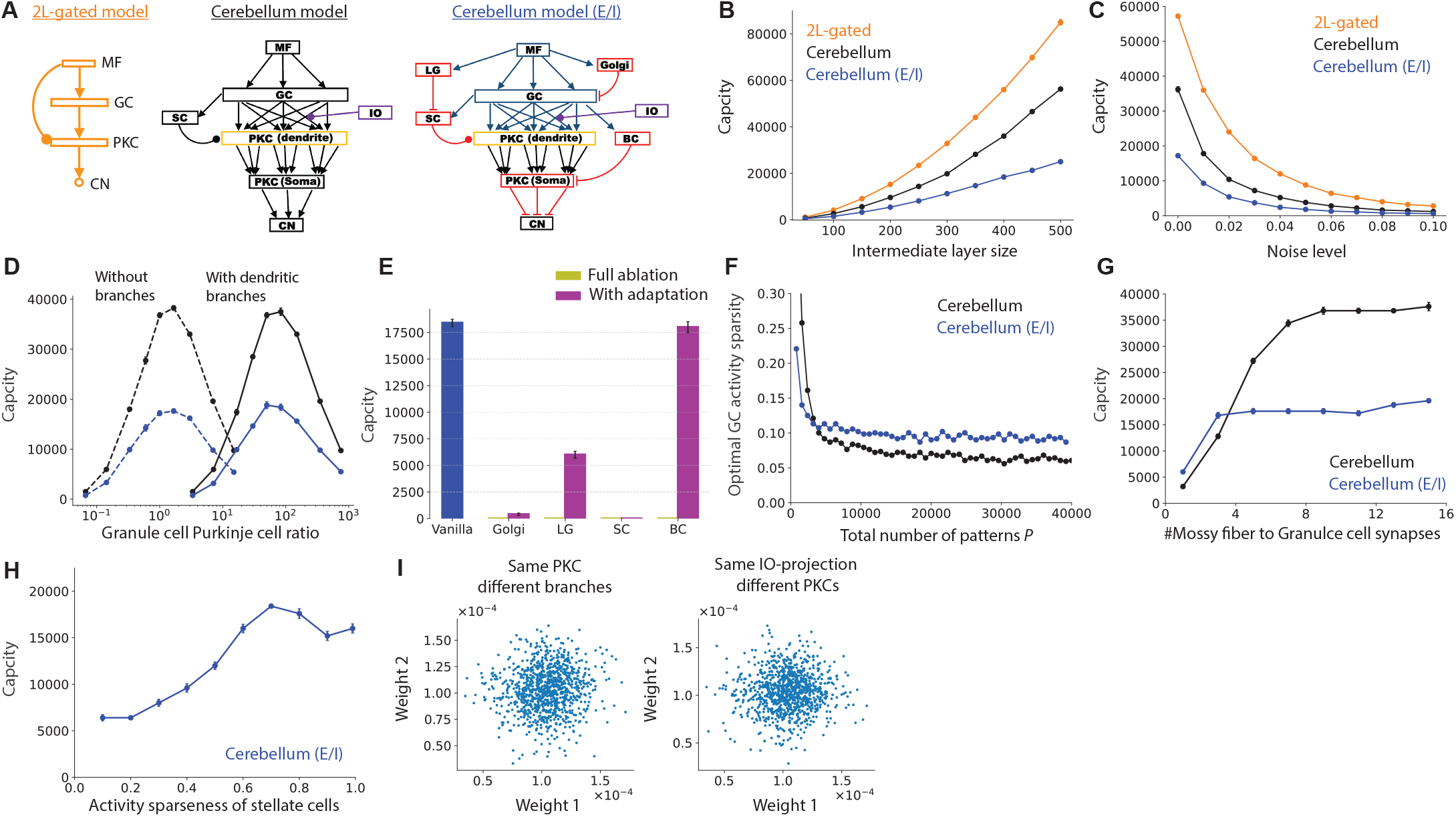
**A)** Illustration of the two-layer gated model and two cerebellar circuit models, without and with E/I constraints. In the cerebellar models, Purkinje cell dendrites and soma are depicted separately to highlight the potentially independent roles of dendritic branches. In the E/I-constrained model, excitatory and inhibitory neurons are shown in blue and red, respectively. Unless otherwise specified, we set the number of mossy fiber (MF) axons to *L*_*MF*_ = 100, granule cells (GC) to *L*_*GC*_ = 400, Golgi cells to *L*_*gol*_ = 50, Lugaro cells (LG) to *L*_*LG*_ = 200, stellate cells (SC) to *L*_*SC*_ = 200, Purkinje cell (PKC) branches to *N*_*branch*_ = 50, Purkinje cells to *L*_*PKC*_ = 8, basket cells (BC) to *L*_*BC*_ = 200, and cerebellar nuclei (NC) neurons to *L*_*NC*_ = 1. See Methods Section 4.2.4 for details. **B)** Capacity of the three models as a function of intermediate-layer size. Capacity is defined as the maximum number of patterns for which the network achieves 90% classification accuracy as in Fig. 1. **C)** Noise tolerance of the three models. The orange curves in panels B and C correspond to those shown in Figs. 1F and I, respectively. **D)** Capacity of cerebellar models with and without dendritic branches (solid vs. dashed lines; *N*_*branch*_ = 50 for solid lines). Black and blue curves denote models without and with E/I constraints, respectively. **E)** Effect of inhibitory neuron ablation on classification capacity. Olive-colored bars (full ablation) are near zero for all cell types. **F)** Optimal granule cell activity sparsity as a function of the number of embedded patterns *P* under a fixed network size (*L*_*GC*_ = 400). Sparsity is defined as the average number of active (non-zero) neurons. **G**,**H)** Capacity under varying sparsity in mossy fiber-to-granule cell connectivity (G) and stellate cell activity (H). **I)** (Left) Comparison of synaptic weights for connections from the same granule cell to the same Purkinje cell but targeting different dendritic branches. (Right) Comparison of synaptic weights for connections from the same granule cell to different Purkinje cells receiving the same inferior olive input.

Nevertheless, the naive mapping contains two clear mismatches with cerebellar anatomy. First, in the model, capacity is maximized when the numbers of granule cells and Purkinje cells are roughly equal (see dashed line in Fig. 5D). This is because, if *L*_*GC*_ and *L*_*PKC*_ denote the numbers of granule cells and Purkinje cells, respectively, then *P* ~ 𝒪 (*L*_*GC*_*L*_*PKC*_), and under a fixed constraint on *L*_*GC*_ +*L*_*PKC*_, capacity is optimized when *L*_*GC*_ ≈ *L*_*PKC*_. By contrast, in the mammalian cerebellum, granule cells outnumber Purkinje cells by more than two orders of magnitude, indicating that the naive model is not well matched to the observed cellular composition. Second, the gating inputs project directly from mossy fibers to Purkinje cells in the naive model, whereas such connections are rare in the real cerebellum [48, 10].

To address these issues and gain further insight into cerebellar circuitry, we modified the model by adding two components (Fig. 5A, middle). First, we introduced Purkinje cell dendritic branches as an intermediate layer between granule cells and the Purkinje cell soma. Purkinje cells have complex dendritic arbors, and individual branches may function as partially independent computational subunits [49, 50]. Recent studies have also demonstrated branch-specific shunting inhibition [51] and plasticity [52], supporting such a computational role. The model naturally accommodates this extension because the PKC-to-CN connections act as a simple summation. Thus, linear summation of branch-specific activity at the soma is sufficient in this framework.

Second, we added a population of stellate cells that receive input from granule cells and project to Purkinje cell dendrites. This makes the gating mechanism more consistent with cerebellar circuitry, because stellate cells receive granule-cell-derived input and project to Purkinje dendrites. Moreover, recent work showed that stellate cell inputs can produce shunting inhibition on Purkinje dendrites, enabling branch-specific gating [51], which is also consistent with the branch-specific computation assumed in the first modification. Note that recent work has emphasized a continuum between stellate and basket cells, which together comprise molecular layer interneurons [10, 53]. Here, we use the term stellate cells to denote the subset of molecular layer interneurons that project primarily to Purkinje dendrites, because dendritic targeting is the key feature required for branch-wise gating. To further align the model with cerebellar circuitry, we also extended it to obey Dale’s law, namely that each neuron is either excitatory or inhibitory (Fig. 5A, right; blue neurons are excitatory and red neurons are inhibitory). As shown below, this extension suggests possible functional roles for several inhibitory neuron classes in the cerebellum [48, 10].

The cerebellum-inspired implementations of the model reduced capacity relative to the original two-layer gated model, but they still achieved supralinear scaling of capacity with intermediate-layer size (black and blue vs. orange in Fig. 5B). In addition, these circuit models retained some robustness to noise, although performance still deteriorated under sufficiently large noise (Fig. 5C).

Introducing dendritic subunits resolved the GC/PKC ratio issue noted above. In the presence of dendritic branches, capacity was optimized when the number of granule cells was much larger than the number of Purkinje cells and approximately matched the total number of Purkinje dendritic branches (i.e., *L*_*GC*_ ≈ *L*_*PKC*_ × *N*_*branch*_; solid vs. dashed lines in Fig. 5D), which is more consistent with cerebellar anatomy. The same qualitative trend was also observed in the E/I-constrained model (blue vs. black lines).

The E/I-constrained model also highlights the potential importance of multiple inhibitory populations in cerebellar circuitry. In addition to Purkinje cells, the model required at least four inhibitory populations to maintain circuit function (see Methods Section 4.2.4 for details). First, a population providing shunting inhibition to Purkinje dendrites was necessary, corresponding to the SC population in Fig. 5A. In addition, three inhibitory populations were required to balance activity at the levels of granule cells, stellate cells, and Purkinje cells. As is well known [6, 54], Golgi cells help regulate excitation-inhibition balance in the granule cell layer. For the remaining two inhibitory populations, which target stellate cells and Purkinje cells, we tentatively map them to Lugaro cells and basket cells, respectively, based on prior anatomical evidence [48]. These assignments, however, are rather provisional, because other inhibitory cell types also target stellate and Purkinje cells, and because the inputs to these populations are not restricted to mossy fibers and granule cells.

We performed ablation analyses of these four inhibitory populations to assess their roles in the model. Complete ablation of any one of the four populations caused a near-complete collapse in performance (olive bars in Fig. 5E; barely visible due to near-zero capacity). By contrast, when the baseline parameters of downstream neurons were allowed to readjust after ablation (see Methods 4.4.5), ablation of Golgi, Lugaro, and stellate cells still caused a marked loss of capacity, whereas the circuit remained comparatively robust to basket cell ablation. Notably, the strong effect of Lugaro cell ablation suggests that disinhibitory input to Purkinje cells may be important for regulating shunting inhibition at Purkinje dendrites, thereby contributing to fast, high-capacity learning in this framework.

Although sparse activity remained important for high capacity, the optimal granule cell activity level was still relatively dense, around ~ 10% (Fig. 5F), because shunting inhibition at dendritic branches introduced an additional source of sparsification in the model. In the model, three to five mossy-fiber inputs per granule cell were sufficient when mossy-fiber-to-granule-cell connections were restricted to be excitatory (Fig. 5G), consistent with previous findings for Marr–Albus-type models [13]. In addition, we found that relatively dense activity in the stellate cell population was favorable for high capacity (Fig. 5H), consistent with the high baseline firing rates reported for these inhibitory neurons [53].

Finally, the model makes an experimentally testable prediction about correlations in synaptic strength. Specifically, among synapses originating from the same granule cell, spine sizes should be uncorrelated across synapses targeting different dendritic branches of the same Purkinje cell, because gating decorrelates synaptic weights (Fig. 5I, left). Similarly, synapses from the same granule cell onto two Purkinje cells receiving input from the same inferior olive neuron should also be uncorrelated (Fig. 5I, right). This prediction differs from that of the traditional Marr–Albus model, which instead predicts correlated synaptic weights in both cases.

## 3 Discussion

Seminal work by Marr, Albus, and Ito postulated the cerebellum as a one-hidden-layer perceptron, in which random nonlinear expansion in the middle layer is followed by supervised learning in the output layer. In this work, inspired by deep learning theory, we propose an alternative view in which the cerebellum operates as a deep neural network with gating. The main motivation for this expansion is that the classification capacity of the network increases quadratically compared with the Marr-Albus model containing the same number of neurons (Fig. 1F). For a network with *L*_*h*_ neurons, this architecture enables classification of *L*_*h*_ times more patterns than the classical model, providing a substantial computational advantage.

We show that, by adding gating to the middle layer, the network can achieve this quadratic capacity using only local three-factor Hebbian plasticity (Fig. 1F). Moreover, the Hebbian updates support fast one-shot learning of patterns, yielding more efficient learning than backprop-based learning in some settings (Figs. 2C and 3D). The proposed model is also capable of solving classification tasks with structured, non-random image inputs (Fig. 3) as well as motor control tasks (Fig. 4).

We further find that the network can be mapped onto a cerebellar microzone circuit containing several inhibitory neuron types (Fig. 5). This mapping makes use of the complex dendritic structure of Purkinje cells, for which previous work has revealed branch-specific shunting inhibition and plasticity. By treating Purkinje cell dendritic branches as basic computational units, the model can retain high capacity while remaining consistent with key anatomical constraints. The model also provides insight into the functional roles of stellate cells (molecular layer interneurons that project to Purkinje cell dendrites) in gating essential for high-capacity local learning, as well as Lugaro cells, which may provide disinhibitory control over Purkinje cells by modulating stellate cell activity. Finally, the model generates experimentally testable predictions. It predicts that, even among Purkinje cells receiving input from the same inferior olive neuron, the spine sizes associated with synapses from the same granule cell should be nearly uncorrelated, because gating by molecular layer interneurons is predicted to control plasticity at granule-cell-to-Purkinje-cell synapses in addition to granule cell activity and inferior olive input. This prediction could be tested using recent EM reconstruction approaches [55, 56].

The proposed model does not capture many known features of the cerebellum. In particular, it is purely feedforward, even though recent work has suggested an important role for recurrent connectivity in cerebellar computation. Nevertheless, this feedforward network was sufficient to solve a reinforcement learning task with spatiotemporal structure. Many other biological features are also omitted from the model, but some, such as neural heterogeneity, would likely affect the performance of the 2L-gated and 1L-vanilla networks in similar ways. The implementations of self-supervised learning in Fig. 3 and the PPO algorithm in Fig. 4 are not biologically plausible, because they require backpropagation in the convolutional and critic components, respectively, even though learning in the 2L-gated and 1L-vanilla circuits was implemented using local plasticity. Both algorithms also require comparison of inputs to a reference signal, the detailed biological implementation of which remains unclear. In the model comparisons, we optimized key parameters, such as activity sparsity and learning rate, individually for each model; however, we did not exhaustively explore the full hyperparameter space. In particular, backprop-based learning could, in principle, achieve learning speeds comparable to the 2L-gated network if the learning rates and initial weights of each layer were finely tuned.

Our work is related to several previous studies. Sezener et al. [51] proposed a cerebellar model that uses gating at Purkinje cell dendrites. Our work builds on this idea by showing that gating enables quadratic classification capacity and rapid learning in a multi-layered setting, thereby revealing a functional advantage of depth in the cerebellum over the classical Marr-Albus model. Walter and and Khodakhah [11] proposed that convergence of Purkinje cell activity onto cerebellar nucleus neurons can implement a denoising operation, thereby improving classification capacity in the presence of neural noise. In our framework, it corresponds most closely to the 2L-vanilla Hebb model in Fig. 2, which, unlike the gated two-layer network, still exhibits only linear capacity with respect to the number of neurons. Muscinelli et al. [57] also examined the functional role of additional depth, but on the input side, in the pontine nuclei layer connecting cortex and cerebellar granule cells. The work revealed a potential role for the pontine bottleneck in capturing low-dimensional latent representations, but it assumed a classical Marr-Albus architecture for the cerebellum itself. A concurrent study by Ruben and Pehlevan [58] focused on heterogeneity in complex-spike-driven LTD at Purkinje cells and showed that such heterogeneity can expand the capacity of a cerebellar microzone, potentially leading to quadratic capacity. However, its applicability beyond the one-shot setting remains unclear because of its reliance on stochasticity. The importance of nonlinear dendritic computation [59, 60] and dendritic gating [61, 62, 63] has also been discussed more broadly in the literature. More generally, many models extending the classical Marr-Albus framework have been proposed to address temporal information processing [64], learning from unsigned supervised signals [65], learning from reward-related information [66], cortico-cerebellar interactions [67], and the functional role of feedback connections [68].

Our work complements these previous cerebellar models in two main ways: first, by showing analytically and numerically that the cerebellum can achieve quadratic pattern classification capacity, rather than the linear capacity of the Marr-Albus model, using biologically plausible one-shot Hebbian plasticity; and second, by demonstrating that the proposed model outperforms the classical cerebellar model in both supervised image classification and motor control tasks, while also largely reproducing the circuit architecture of a cerebellar microzone with several cell types under excitatory/inhibitory constraints. More broadly, the faster learning curves relative to backprop that we observed in Figs. 2C and 3D support the hypothesis that the specific neural architecture of the brain promotes efficient learning. In this view, the multilayer feedforward organization of the cerebellum may not simply reflect anatomical complexity, but may instead constitute an evolutionary solution for combining rapid learning with high representational capacity.

## 4 Methods

### 4.1 Task setting

#### 4.1.1 Perceptron problem

Following previous computational studies of the cerebellum [6, 7, 69, 13], we considered a binary classification task on input patterns. We constructed *L*_0_-dimensional input patterns 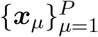 by independently sampling each element as

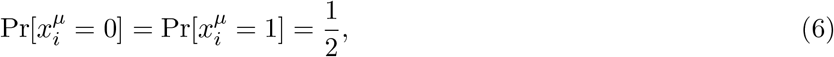

for *i* = 1, …, *L*_0_ and *µ* = 1, …, *P*. For each pattern ***x***_*µ*_, we randomly assigned a binary label 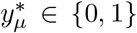 according to

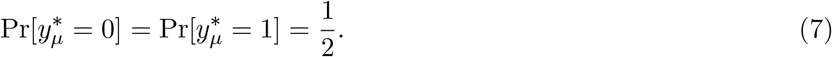

The task is to infer the label 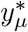 from the input ***x***_*µ*_.

For pattern classification from noisy inputs, we generated a noisy input 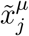 by independently flipping each input element with probability *q*:

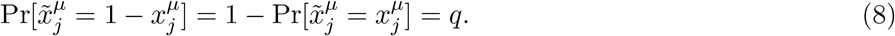

#### 4.1.2 Omniglot character classification

In the Omniglot task shown in Fig. 3, we first randomly divided the 1,623 characters included in the Omniglot dataset[39] into 599 pre-training characters, which were used to train the convolutional neural network (CNN), and 1,024 characters, which were used to train the subsequent feedforward networks. For each of the 1,024 characters, we selected 15 images for training and 2 images for testing. Accordingly, memorization performance was defined as the average classification accuracy on the 15 × 1024 training images, whereas generalization performance was defined as the classification accuracy on the 2 × 1024 test images. Binary labels were assigned randomly to characters, irrespective of the writing system to which they belonged. We used this binary classification setting as a natural extension of the perceptron task to naturalistic images. We compressed the original 105 × 105 pixel images to 52 × 52 pixels for memory efficiency.

The CNN was pre-trained using noise contrastive estimation (NCE) learning [42, 43]. We used a simple CNN consisting of two 3 × 3 convolutional layers (with 32 and 64 channels, respectively), each followed by average pooling, and a final 128-dimensional dense layer. For each image in the pre-training dataset, we randomly selected another image of the same character and used it as the positive sample for NCE. At each weight update, we sampled a mini-batch of 100 images and used those images as the pool of negative samples. Thus, each sample had 1 positive example, with which the final-layer representation should be similar, and 99 negative examples, from which the representation should be distinct. We trained the network for 10 epochs using the Adam optimizer with learning rate 10^−3^.

Using this pre-trained network, we obtained a 128-dimensional representation for each training image and used these representations as inputs to the 2L-gated, 1L-vanilla, and 2L-vanilla networks. Please see the repository linked in the Code Availability section for further details.

#### 4.1.3 Reinforcement learning task

For the reinforcement learning tasks implemented in Fig. 4, we used `reacher_v5` and `pendulum_v1` from the Gymnasium library [44]. Each trial lasted 80 steps in `reacher_v5` and 192 steps in `pendulum_v1`. The reward in `reacher_v5` was defined as

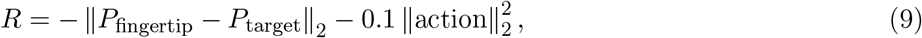

where *P*_fingertip_ and *P*_target_ are the two-dimensional coordinates of the robot arm fingertip and target, respectively, and `action` is the torque vector applied to the robot arm. In `pendulum_v1`, the reward was defined as

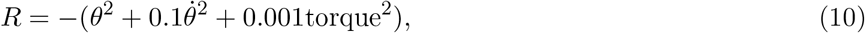

where *θ* is the pendulum angle. The maximum achievable reward was zero in both tasks. Please see Ref. [44] for further details of these tasks.

### 4.2 Model configurations

#### 4.2.1 Two-layered gated model (2L-gated)

##### Dynamics

We constructed a feedforward network with two intermediate layers and gating as below. We denote the number of neurons in the *k*-th layer by *L*_*k*_ (i.e., *L*_0_ for the input layer, *L*_1_ for the first intermediate layer, and so on). The activity of the first intermediate layer is given by

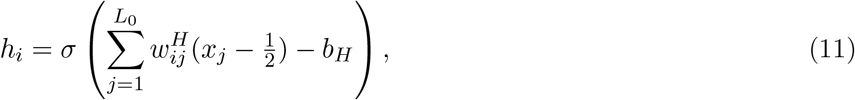

for *i* = 1, …, *L*_1_, where 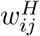 is the synaptic weight from neuron *j* to neuron *i, b*_*H*_ is the baseline membrane bias, and *σ*(*u*) is the step function, which returns 1 if *u >* 0 and 0 otherwise. The activity of the second intermediate layer is defined as

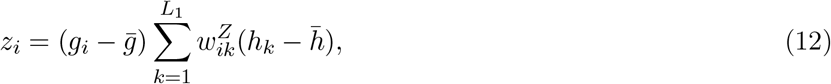

for *i* = 1, …, *L*_2_, where 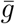 and 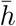 are the baseline activities, and *g*_*i*_ is an input-dependent gating unit that modulates the activity of neurons in the second intermediate layer. We set the gating unit activity as

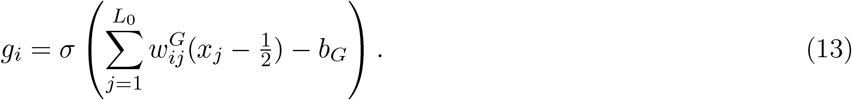

Finally, summing over the second-layer activity yields the output *y*:

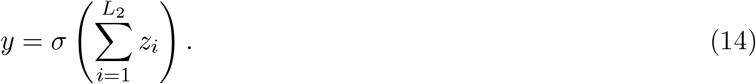

In the cerebellar interpretation of the model, *x*_*i*_ corresponds to a mossy fiber, *h*_*i*_ to a granule cell, *z*_*i*_ to a dendritic compartment of a Purkinje cell, and *y* to a cerebellar nuclei neuron. A more biologically grounded implementation is described in Method 4.2.4.

##### Weight configuration

We sampled each element of 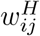 and 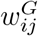 independently from a normal distribution:

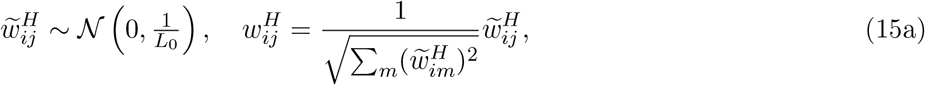

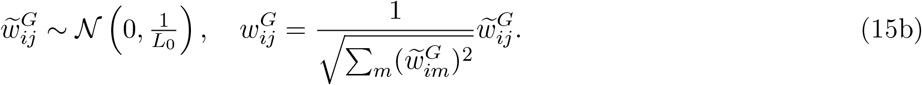

Here, we introduced postsynaptic weight normalization, which reduces neuron-to-neuron variability in activity sparseness in the sparse-activity limit. In models with sparse connectivity, we instead generated *w*^*H*^ using connectivity sparsity *ρ*_*H*_ as

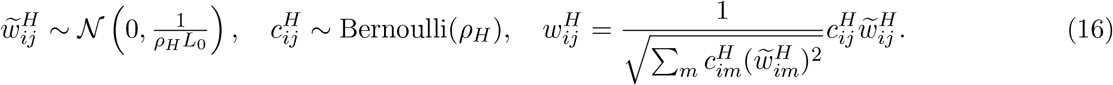

Sparse *w*^*G*^ was generated in the same manner. The baseline biases *b*_*H*_ and *b*_*G*_ were set to constant values that were optimized for capacity (Fig. 1D).

##### Three factor Hebbian weight update

The connections between the two intermediate layers, {*h*_*i*_} and {*z*_*i*_}, were constructed by a three-factor Hebbian rule modulated by the presynaptic input *h*_*j*_, the gating signal *g*_*i*_, and the teaching signal 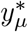:

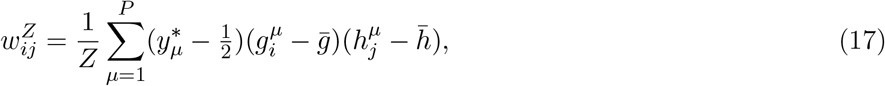

where 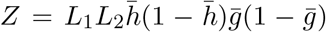 is the normalization factor, and 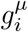 and 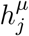 are the activities of *g*_*i*_ and *h*_*j*_, respectively, for input ***x*** = ***x***_*µ*_. Although this is not the optimal weight configuration, it is biologically plausible as it only requires information locally available to synapse 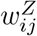. Moreover, learning can be achieved in a one-shot manner, meaning that only a single exposure to each pattern is required. This construction can also be written in an online sample-by-sample form, where after presentation of the *p*-th pattern the weight becomes

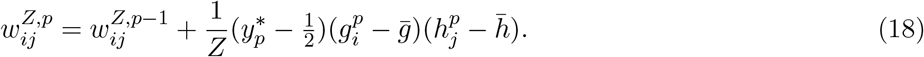

##### Error-based weight update

Given the *µ*-th pattern ***x***_*µ*_, the error-based iterative weight update was implemented as

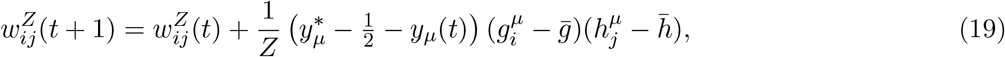

where

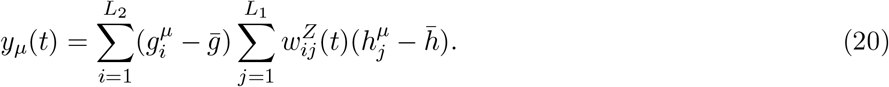

In this iterative update rule, the teaching signal is the prediction error, 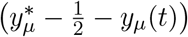, rather than the target signal 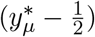 used in the three factor Hebbian learning.

#### 4.2.2 One-layer model with random expansion (1L-vanilla)

We next consider a standard one-hidden-layer network with random expansion in the intermediate layer. This model has been studied extensively in both neuroscience [30, 32] and machine learning literature [70]. We define the binary firing activity of intermediate-layer neuron *h*_*i*_ as

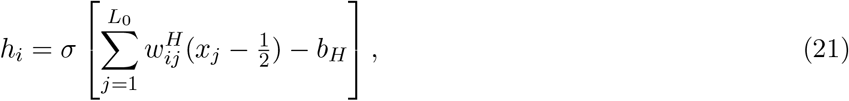

where *b*_*H*_ is the bias term, and the weights 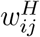 are L2-normalized Gaussian random variables:

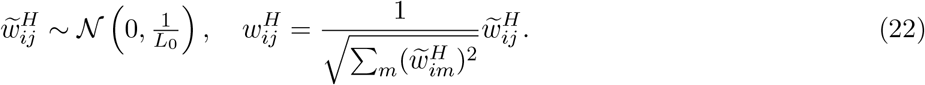

The output *y* is then given by

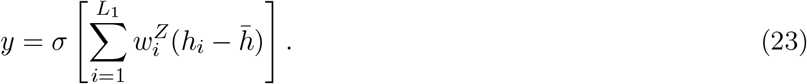

Here, 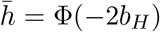 is the mean activity of *h*_*i*_, and *w*^*Z*^ is set according to an association of the teaching signal and presynaptic activity:

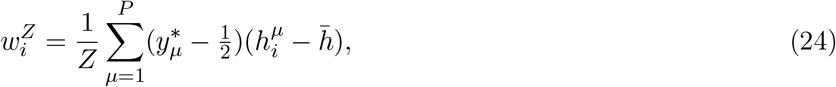

with 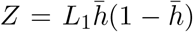. As before, 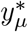 denotes the teaching signal and 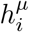 is the intermediate-layer activity for ***x*** = ***x***_*µ*_.

#### 4.2.3 Two- and three-layered vanilla models

##### Two-layered vanilla model with Hebbian-type learning

The two-layered model without gating (i.e., 2L-vanilla) used for Hebbian-type learning (depicted in Fig. 2B) was defined as

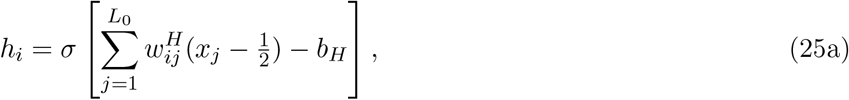

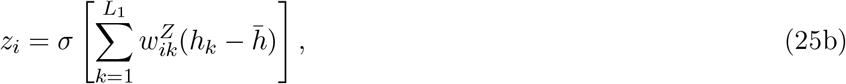

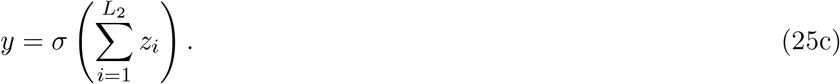

As before, *w*^*H*^ was initialized as L2-normalized Gaussian random weights. A naive extension of the one-hidden-layer model above yields the following associative construction for *w*^*Z*^:

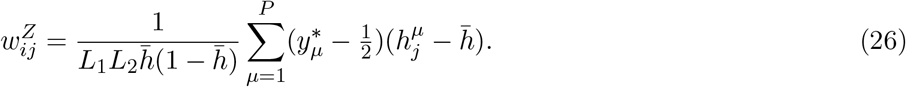

##### Two-layered vanilla model with backprop

For backprop training, we used a slightly different network architecture and weight initialization, because the network needed to be differentiable. Specifically, we implemented the two-layered vanilla model as

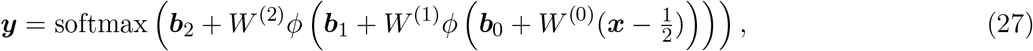

where *ϕ*(*x*) is the element-wise rectified linear unit. Unlike the previous networks, which had a single output, here we set ***y*** = [*y*_0_, *y*_1_] to be a two-dimensional output vector and used the softmax function to generate the output probabilities, to mimic the standard machine learning setting and accelerate the learning process.

##### Three-layered vanilla model with backprop

Similarly, the three-layered network was defined as

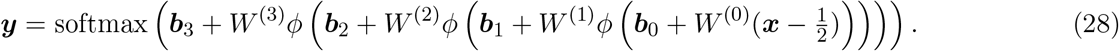

In both two-layered and three-layered networks, all weights were trained using the Adam optimizer, following common practice in deep neural network training. We initialized the weights *W*^(*ℓ*)^ as Gaussian random variables with mean zero and variance 1*/L*_*k*_ (without L2 regularization). The bias parameters ***b***_*k*_ were initialized to zero.

#### 4.2.4 Cerebellum model

We considered two implementations of the 2L-gated model in cerebellar circuitry. The first is a minimal extension that does not impose Dale’s law, whereas the second additionally enforces excitatory/inhibitory (E/I) constraints.

**Table 1:** Acronyms of cell types.

| Acronym | Name |
| --- | --- |
| MF | Mossy fiber |
| GC | Granule cell |
| PKC | Purkinje cell |
| PKD | Purkinje cell dendrite |
| PKS | Purkinje cell soma |
| CN | Deep cerebellar nuclei |
| IO | Inferior olive |
| SC | Stellate cell |
| Golgi | Golgi cell |
| LG | Lugaro cell |
| BC | Basket cell |

##### Cerebellum model without Dale’s law

In this implementation, we introduced two modifications to the 2L-gated model to better match cerebellar anatomy. First, the gating signal was assumed to arise from the granule cell layer through stellate cells, rather than being projected directly from the mossy fiber layer as in the original 2L-gated model. We introduced this modification because there is no clear direct gating input from mossy fibers to Purkinje cells. Second, we separated Purkinje cells into dendritic and somatic compartments, assuming branch-specific gating and linear dendritic summation. This extension increases the capacity of the circuit while approximately capturing dendritic computation in Purkinje cells.

We constructed the network as follows. The input layer corresponds to mossy fiber (MF) activity, denoted by *r*^*MF*^. MF activity projects to the granule cell (GC) layer according to

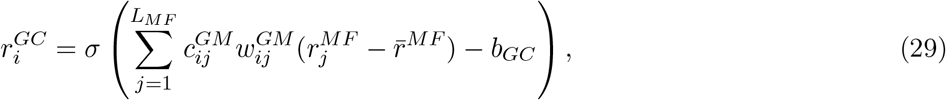

where 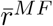 is the mean MF activity and *b*_*GC*_ is the GC baseline that controls activity sparsity. Here, 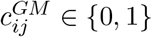 denotes the connectivity between MF *j* and GC *i*, while 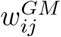 is the synaptic weight when 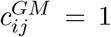. The connectivity variable 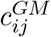 was introduced to capture sparse MF-to-GC connectivity.

GC activity projects both directly to Purkinje cell dendrites (PKD) and indirectly through stellate cells (SC):

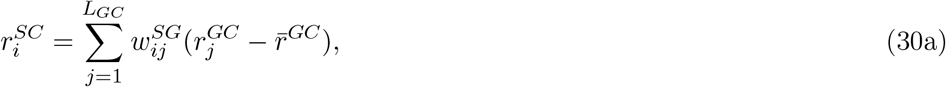

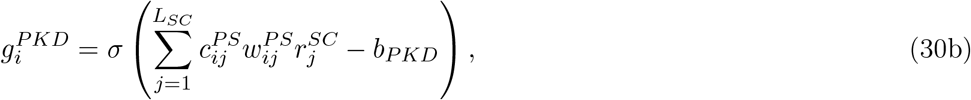

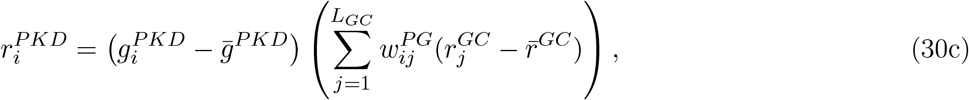

where 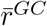 is the mean GC activity, and weights *w*^*SG*^, *w*^*PS*^, *w*^*PG*^ represent GC-to-SC, SC-to-PKD, GC-to-PKD weights, respectively. Here, 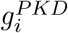 represents branch-specific gating at the *i*-th dendritic branch.

The somatic activity of each Purkinje cell was taken to be the sum of the activities of its dendritic branches. Denoting the set of dendrites belonging to the *i*-th Purkinje cell by Ω_*i*_, we have

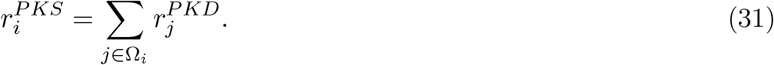

Finally, the deep cerebellar nuclei (CN) neuron pools activity over local Purkinje cells:

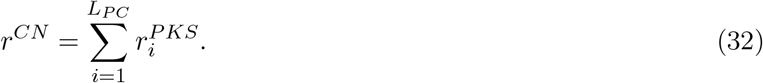

We generated the connectivity from MF to GC, 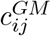, randomly while constraining the in-degree of each GC to *K*_*GM*_ connections. Here, *GM* denotes the projection from mossy fibers to granule cells, following the matrix notation. As before, we normalized the total input weight so that the L2 norm remained constant regardless of the number of projections *K*_*GM*_ :

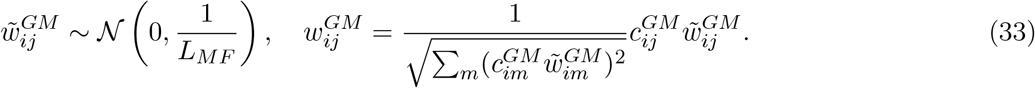

As in the MF-to-GC projection, we constrained SC-to-Purkinje dendrite connections to be sparse, because each SC is expected to project to only a small number of Purkinje dendrites. The connectivity 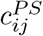 was generated randomly with in-degree constrained to *K*_*PS*_, and 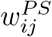 was sampled randomly from a normal distribution with L2 normalization. The GC-to-SC weights 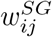 were generated in the same manner, but with full connectivity.

The GC-to-Purkinje dendrite weights 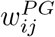 were the locus of plasticity in the model, as before. Using the GC neural activity and dendritic gating activity under pattern *µ*, denoted 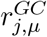 and 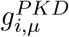, we set the weight as

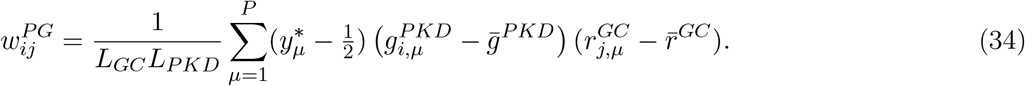

##### Cerebellum model with Dale’s law

We next constructed a version of the model that approximately satisfies Dale’s law by introducing additional inhibitory interneurons.

To impose Dale’s law on MF-to-GC connectivity, we introduced Golgi cells, which receive input from MF and inhibit GC. We defined the activities of GC and Golgi cells as

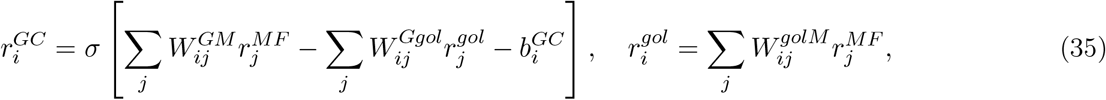

where *W*^*GM*^, *W*^*Ggol*^, and *W*^*golM*^ denote MF-to-GC, Golgi-to-GC, and MF-to-Golgi connections, respectively.

We constructed the weights *W*^*GM*^, *W*^*Ggol*^, and *W*^*golM*^ using non-negative matrix factorization (NMF). We first generated a weight matrix 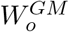 as a sparse random matrix in which each postsynaptic neuron received input from only 2*N*_*syn*_ mossy fibers, without imposing the E/I constraint. We then separated this weight into excitatory and inhibitory components by setting 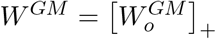 and 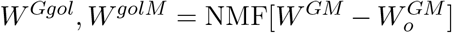, where [*x*]_+_ ≡ max(0, *x*) is applied element-wise and NMF returns two non-negative matrices *A* and *B* such that *AB* ≈ *C* for a given non-negative matrix *C*. The baseline parameter 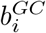 was then set as

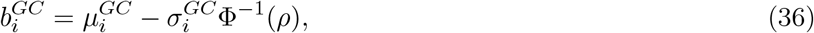

where *ρ* is the target activity sparsity, and 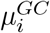 and 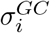 are the mean and standard deviation of 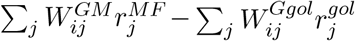 over input patterns. Although this construction is not strictly biologically plausible, it could in principle be approximated through homeostatic plasticity.

To impose Dale’s law on the SC pathway, we introduced Lugaro cells (LG), which can provide disinhibitory input to SC, as well as to basket and Golgi cells. LG receive major input from Purkinje cells in addition to mossy fiber input, although we did not explicitly model the former here. The activities of SC and LG were defined as

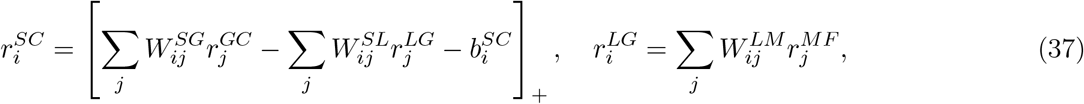

where [*x*]_+_ ≡ max(0, *x*), and *W*^*SG*^, *W*^*SL*^, and *W*^*LM*^ denote GC-to-SC, LG-to-SC, and MF-to-LG connections, respectively.

We generated *W*^*SG*^ as a sparse random non-negative matrix in which each SC received input from only 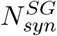 GCs. Similarly, we sampled the product *W*^*SL*^*W*^*LM*^ as a sparse random non-negative matrix with in-degree 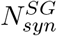, and then generated *W*^*SL*^ and *W*^*LM*^ using NMF. We subsequently scaled *W*^*SL*^ by

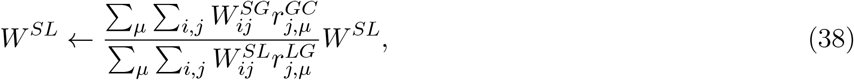

to enforce approximate E/I balance. The baseline 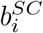 was set as

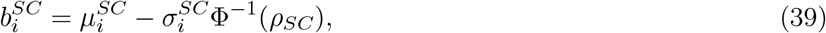

where 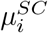 and 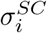 are the mean and standard deviation of the *i*-th SC activity over patterns.

Given GC and SC activity, the activity of Purkinje cell dendrites was defined as

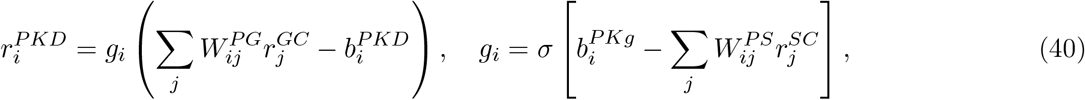

where *W*^*PG*^ and *W*^*PS*^ denote the GC-to-Purkinje dendrite and SC-to-Purkinje dendrite connections, respectively. Because *g*_*i*_ ∈ {0, 1}, the activity of the *i*-th dendrite is shunted to zero when SC input is larger than the threshold for shunting inhibition 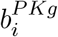. If not, inputs from GC are propagated to the PKC soma. The SC-to-Purkinje gating weights were generated randomly under a sparse connectivity constraint, and 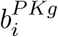 was set as

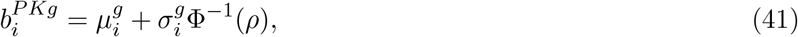

so that, on average, only a fraction *ρ* of dendrites were shunted.

We set the GC-to-dendrite weights of Purkinje cells as

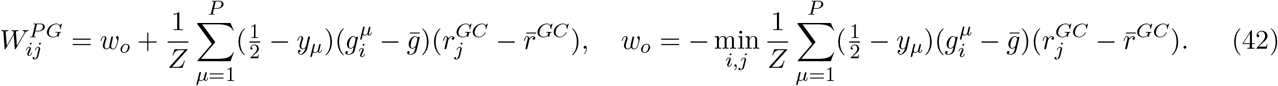

We defined 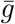 and 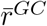 as the mean activity levels of the gating signal and excitatory input over patterns and dendritic branches, respectively. The normalization factor was *Z* = *L*_*GC*_*L*_*PKD*_. By construction, 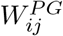 is non-negative. The baseline 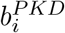 was set to

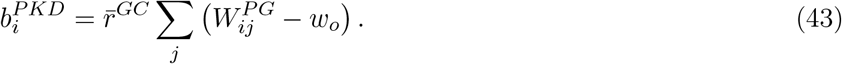

The activity of the Purkinje cell soma was then defined as a linear sum over dendritic branches, together with inhibition from basket cells (BC) and a global inhibitory term:

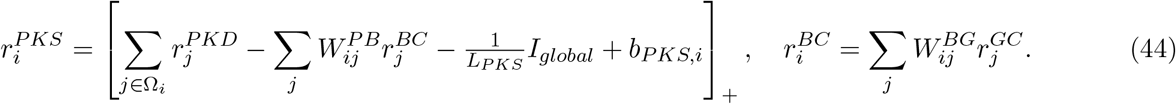

This expression arises from decomposing the original 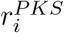 expression in the previous section into positive and negative components. The BC-to-Purkinje weights *W*^*PB*^ and GC-to-BC weights *W*^*BG*^ were defined such that the product *W*^*PB*^*W*^*BG*^ satisfied

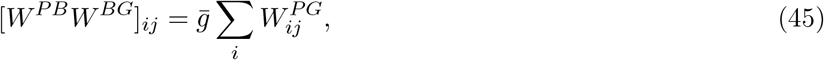

using non-negative matrix factorization. Unlike *W*^*Ggol*^, *W*^*golM*^, *W*^*SL*^, and *W*^*LM*^, the construction of *W*^*PB*^ and *W*^*BG*^ depends on the learned patterns and therefore requires adaptation, although the summation over *i* makes this term nearly constant. The global inhibition term was defined as

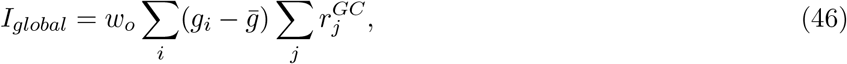

and although its exact biological implementation is unclear, such a signal could in principle arise through several mechanisms, including astrocyte-mediated effects. Finally, the baseline term was given by

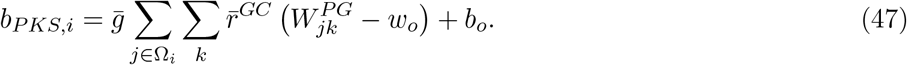

The offset *b*_*o*_ was defined as 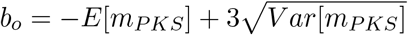, where

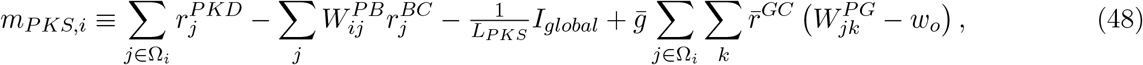

so as to minimize truncation of 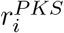. We computed the mean and variance over both neurons *i* = 1, …, *L*_*PKS*_ and patterns *µ* = 1, …, *P*.

The CN activity pools Purkinje cell outputs via:

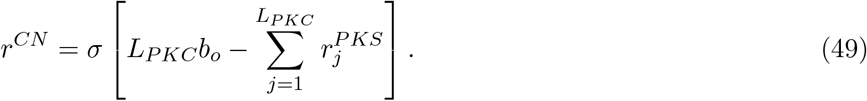

This is only one possible mapping of the proposed 2L-gated network onto cerebellar circuitry, and other implementations are clearly possible.

### 4.3 Analysis of classification capacity

#### 4.3.1 Analysis of the intermediate layer activity

We estimate the classification performance by calculating the signal-to-noise ratio. To this end, we first analyze the statistics of the intermediate layer activity at *L*_0_, *L*_1_, *L*_2_, *P* ≫ 1 limit. For the ease of notation, we denote the membrane variables of the first intermediate layer neurons and gating neurons under ***x*** = ***x***_*µ*_ by

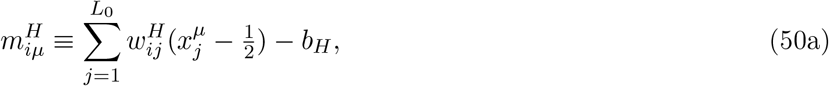

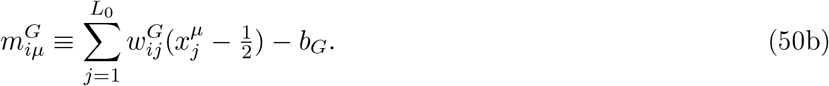

Taking the expectation over random binary patterns {***x***_*µ*_}, we have

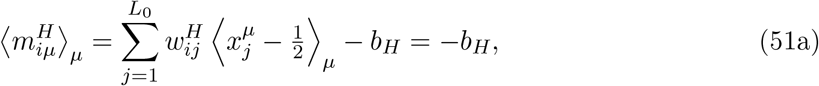

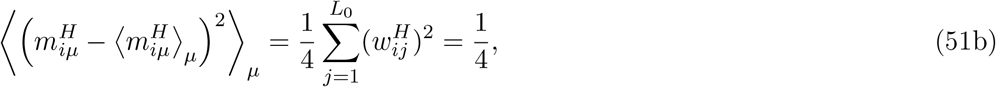

where ⟨·⟩_*µ*_ denotes the average over random patterns. The second equation follows from the weight normalization in Eq. 15a. Therefore, the membrane potential variable 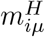 approximately follows 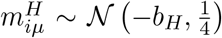. This means that the firing probability of the first intermediate layer neuron *h*_*i*_ follows

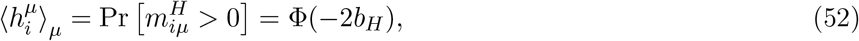

where

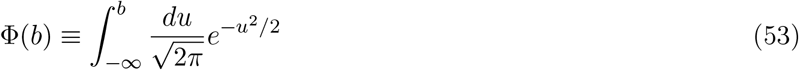

is the cumulative distribution function of the standard Gaussian. Hence, the baseline activity 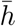 in Eq. 12 should be set to

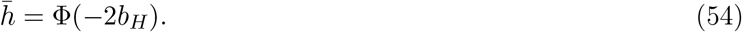

From the symmetry between 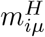 and 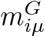, we also have 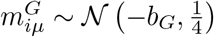 and 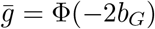.

When the total number of patterns *P* is much larger than the input layer size *L*_0_, correlation between patterns are not negligible. For two patterns *µ* ≠ *ν*, taking the average over intermediate layer neurons, for *Q* = *H* or *G*, we have

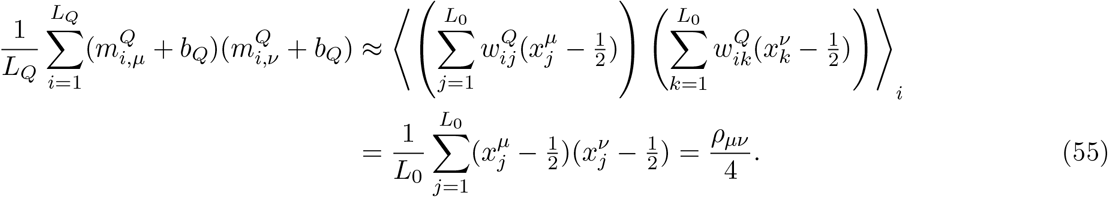

In the last line, we defined the pattern correlation at the input layer by

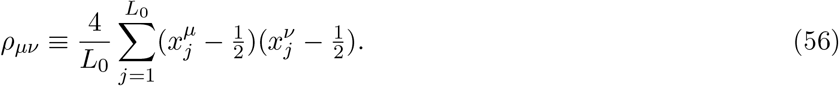

Due to the nonlinear transformation by *σ*(·), it is difficult to evaluate how the input correlation 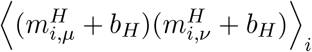 is translated into the output correlation 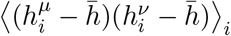 exactly. However, using 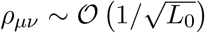, we can evaluate the output correlation up to the second order term with respect to *ρ*_*µν*_ with Taylor expansion [33]. The joint probability of 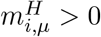 and 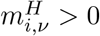 is written as

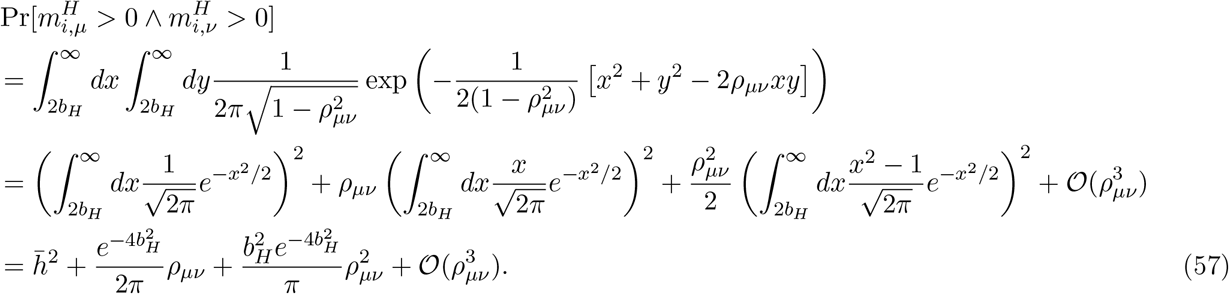

Therefore, taking the expectation over neurons,

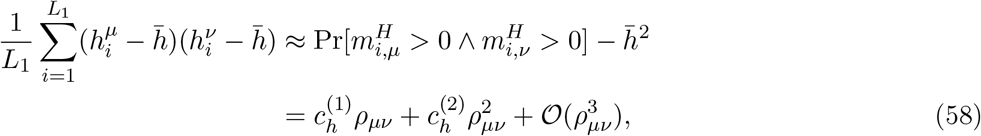

where

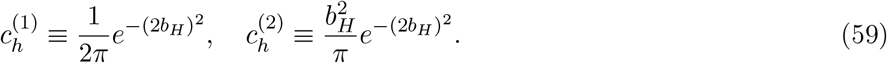

The first line of Eq. 58 is asymptotically true at a large *L*_1_ limit. Thus, the correlation between patterns are scaled by the factor 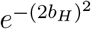 in the intermediate layer activity, indicating that when the bias *b*_H_ is large (i.e., when the intermediate layer activity is sparse), the correlation is suppressed exponentially. For brevity, we define the average correlation in the first intermediate layer activity under patterns *µ* and 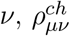, by

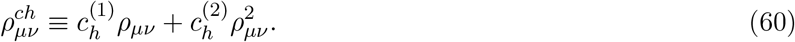

From a parallel calculation, we have

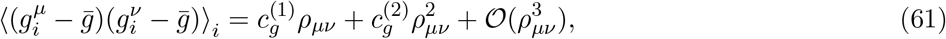

where 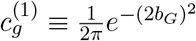 and 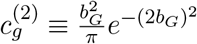.

#### 4.3.2 Signal-to-noise ratio analysis of two-layer gated network

Given input ***x*** = ***x***_*µ*_, the output membrane variable 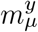 is decomposed into the signal and noise components as below:

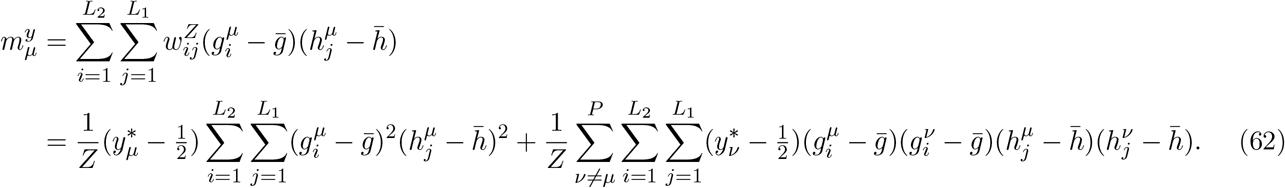

Below, we evaluate the amplitude of the signal and noise to estimate the signal-to-noise ratio as a function of the activity sparseness. Fully analytical setting is studied in Sec. 4.3.4.

Given random ***w***^*H*^ and ***w***^*G*^, the numbers of active neurons in the first intermediate layer and the gating units are on average uncorrelated. Thus, the probability of having *m* active neurons in the first intermediate layer and *n* active neurons in the gating units, is approximately written as

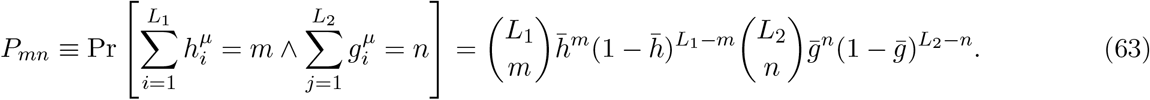

Under this condition, the signal component follows

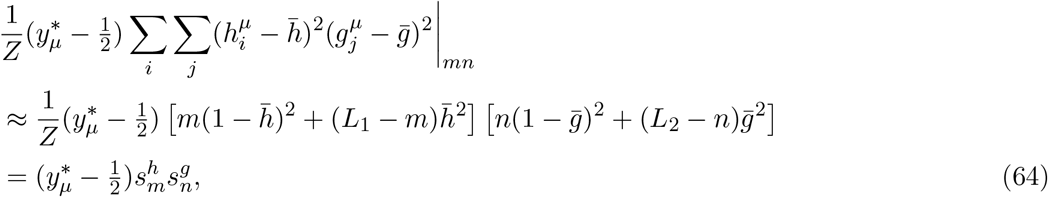

where |_*mn*_ indicates the condition, 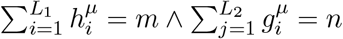, and 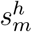 and 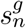 are defined by

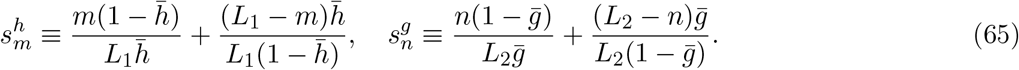

On the other hand, after a relatively straightforward calculation (see Section 4.3.5), the variance term follows:

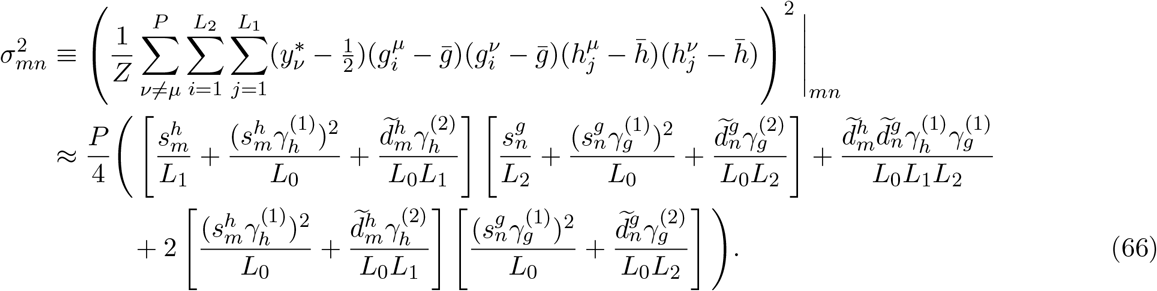

Here, for brevity, we introduced following variables:

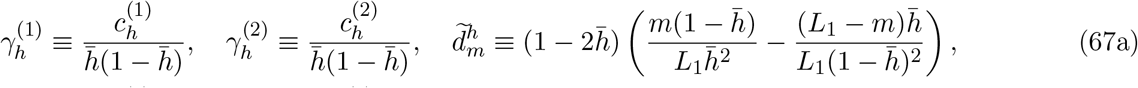

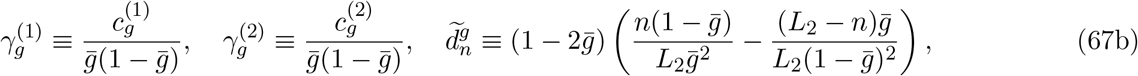

and used 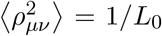 and 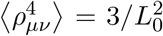. Therefore, from Eqs. 63, 64, and 92, the probability of a correct classification becomes

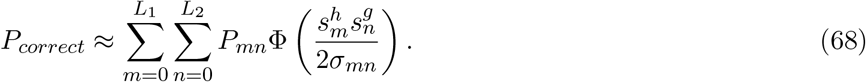

As we discuss in Sec. 4.3.4, the signal terms 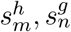 are 𝒪(1), while the noise term *σ*_mn_ scales with 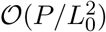 in general. However, *γ* terms decays exponentially as a function of activity bias *b*, enabling 𝒪(*P/*(*L*_1_*L*_2_)) scaling at the sparse limit.

#### 4.3.3 One-intermediate layer model with random expansion

For convenience, let us denote the membrane variables under ***x*** = ***x***_*µ*_ by

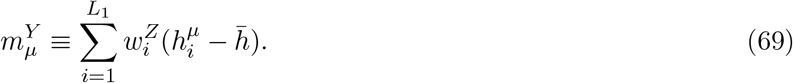

As before, we can estimate the probability of correct classification by evaluating the signal-to-noise ratio. Decomposing 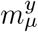 into the signal and noise components:

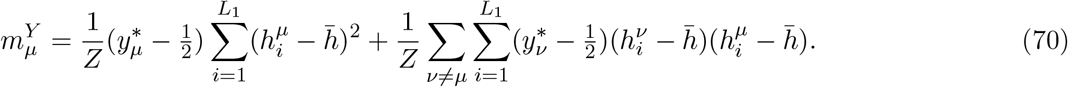

Because both the signal and noise scales with the number of active units, ∑_*i*_ *h*_*i*_, we evaluate the ratio for each activity level *k* = 0, 1, …, *L*_1_. The probability of having *k* active neurons in the intermediate layer approximately follows

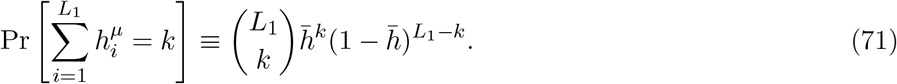

In this condition, the signal component of Eq. 70 becomes

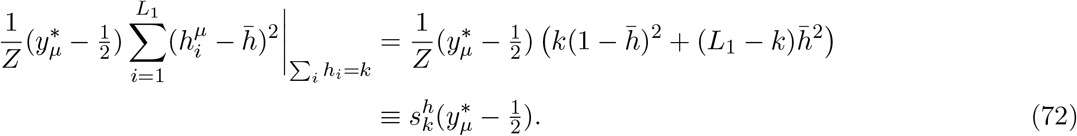

where 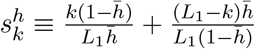. On the other hand, the noise term follows

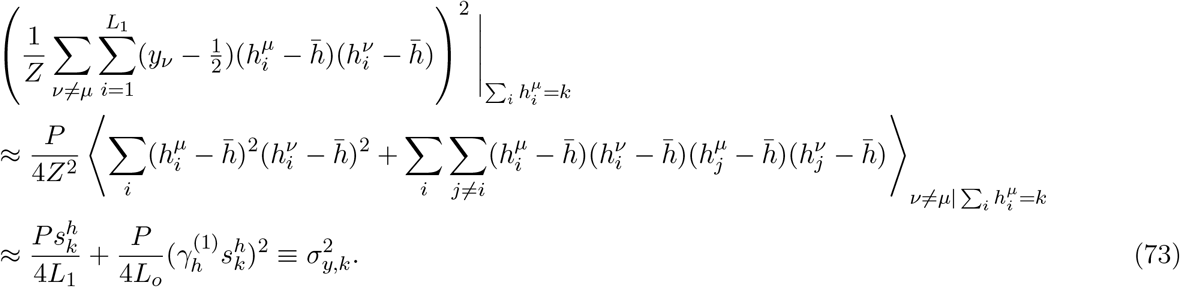

In the last line, we used

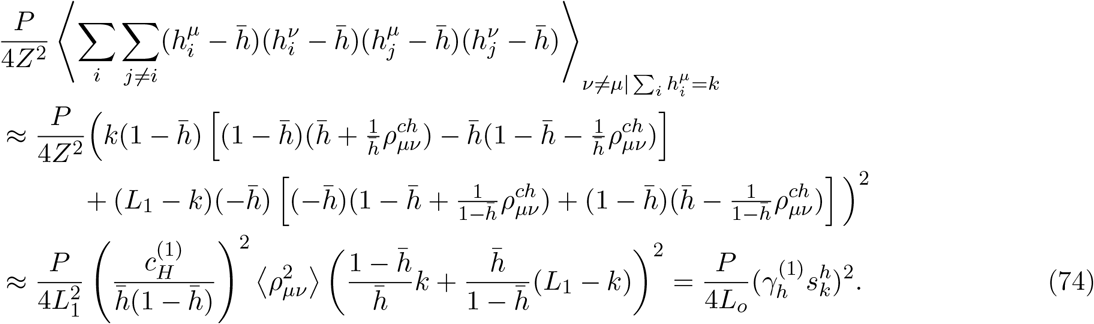

Although the second term of the noise (Eq. 73) scales with *P/L*_0_, because 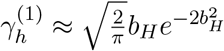. this term becomes negligible under a large bias *b*_*H*_. In this regime, *P* ≲ *L*_1_ is sufficient for achieving large signal-to-noise ratio, as suggested previously [30]. Then, the probability of a correct classification becomes

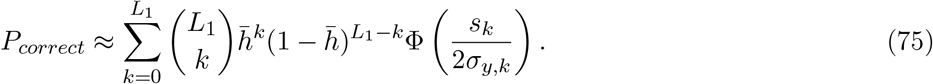

#### 4.3.4 Asymptotic behavior of the two-hidden-layer gated network

Let us consider the limit where 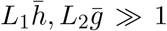 while 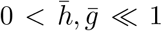. From the central limit theorem, 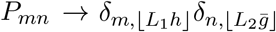, where *δ*_*m,n*_ is the Dirac delta and ⌊*x*⌋ is the floor function. At *m* = *L*_1_*h* and 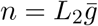, we have 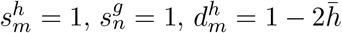 and 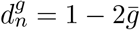. Thus, the probability of a correct classification obeys

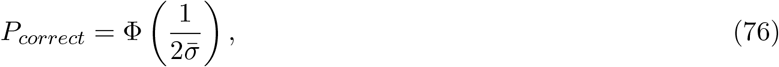

where

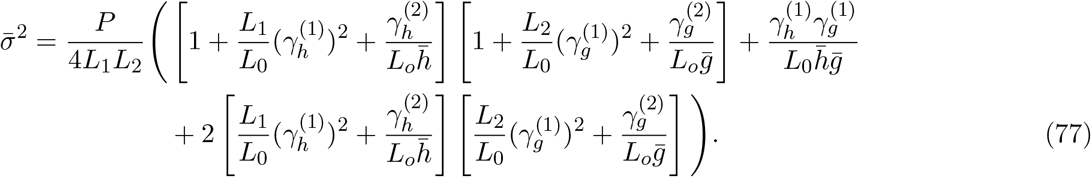

Therefore, the signal-to-noise ratio is approximately written as,

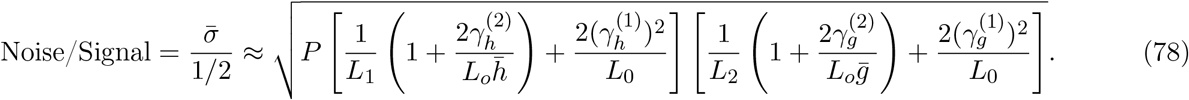

Defining

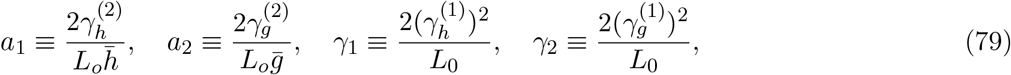

we get Eq. 3 in the main text. As shown in Fig. 6A, 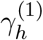 and 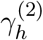 terms decays in sub-Gaussian fashion as a function of the baseline *b*_*H*_.

**Figure 6:**
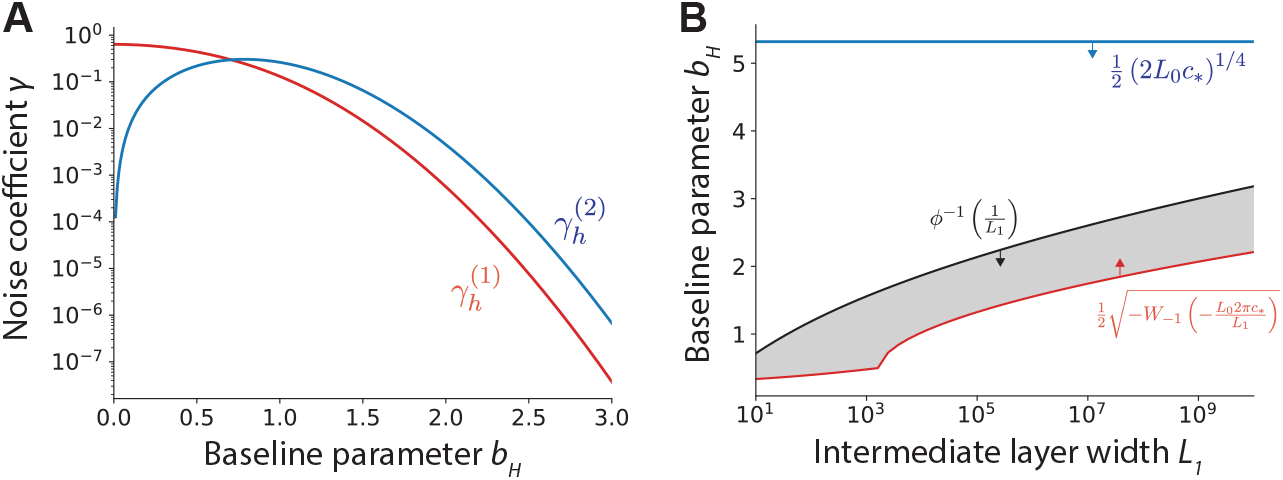
**A)** Noise coefficients 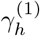 and 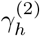 (Eqs. 93a) as a function of baseline *b*. **B)** Upper and lower bounds on the baseline parameter *b*_*h*_ under *L*_0_ = 100 and *c*_∗_ = 0.1. The shaded area achieves quadratic scaling under our approximated theory.

Let us further assume that *L*_1_ = *L*_2_ and *b*_*h*_ = *b*_*g*_. Then, in order to achieve 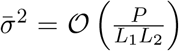, *L*_0_-dependent terms in the equation above need to satisfy

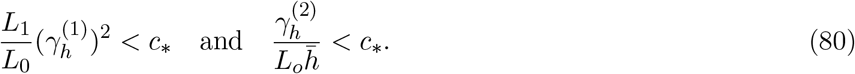

where *c*_∗_ is a small positive constant. Substituting 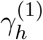 and 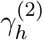 with their definitions (Eqs. 93a and 59), the inequalities above are rewritten as

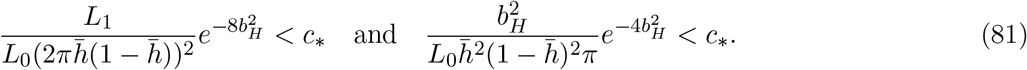

When the bias *b*_*H*_ is large, using an asymptotic expression of the cumulative Gaussian distribution 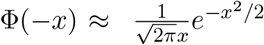, we get 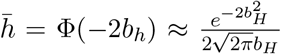. Substituting 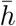 with this approximation, the equations above are rewritten as

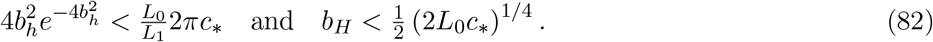

Using Lambert W functions *W*_0_ and *W*_−1_, the first inequality is rewritten as 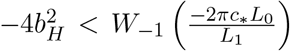. The inequality is also satisfied if 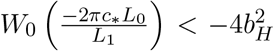, but this region violates the assumption that *b*_*H*_ is large.

Because *W*_−1_(*x*) ≤ −1 for 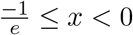, the inequality above is rewritten as 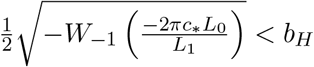.

Moreover, considering finite size effect, the activity sparsity needs to satisfy 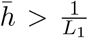. Denoting 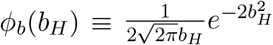, needs to satisfy 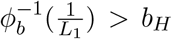, Therefore, to satisfy 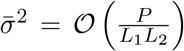, the bias *b*_*H*_ needs to satisfy

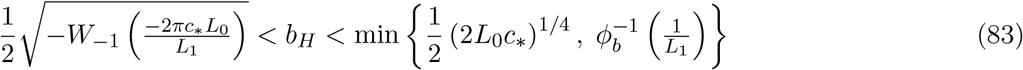

Fig. 6B captures these lower and upper bounds for various *L*_1_ under *c*_∗_ = 0.1. When *L*_0_ is large (*L*_0_ ≫ 1), we have 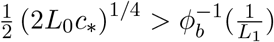, but the lower bound for *b* for sufficient sparsity is smaller than this upper bound robustly across *L*_1_ (red vs. black lines in Fig. 6B; shaded area represents *b*_*H*_ satisfying Eq. 83), suggesting a quadratic scaling over a wide range of *L*_1_.

#### 4.3.5 Estimation of the noise term in the signal-to-noise ratio analysis of two-hidden-layer gated networks

The noise term of the output membrane potential (Eq. 62) is given as

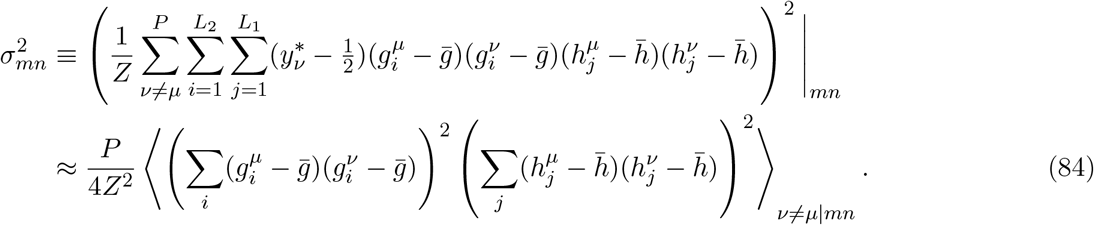

The *h*-dependent term is decomposed into

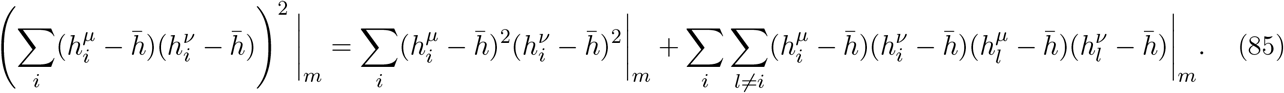

Because patterns typically have non-zero correlation with each other, we have

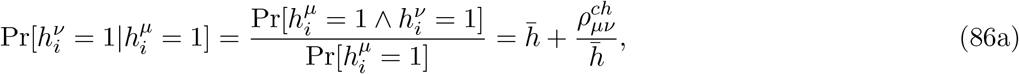

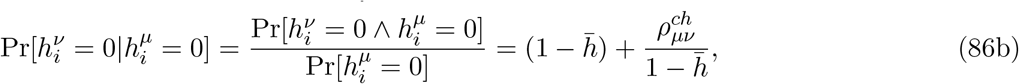

where 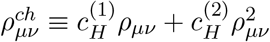. Thus, the first term of Eq. 85 becomes

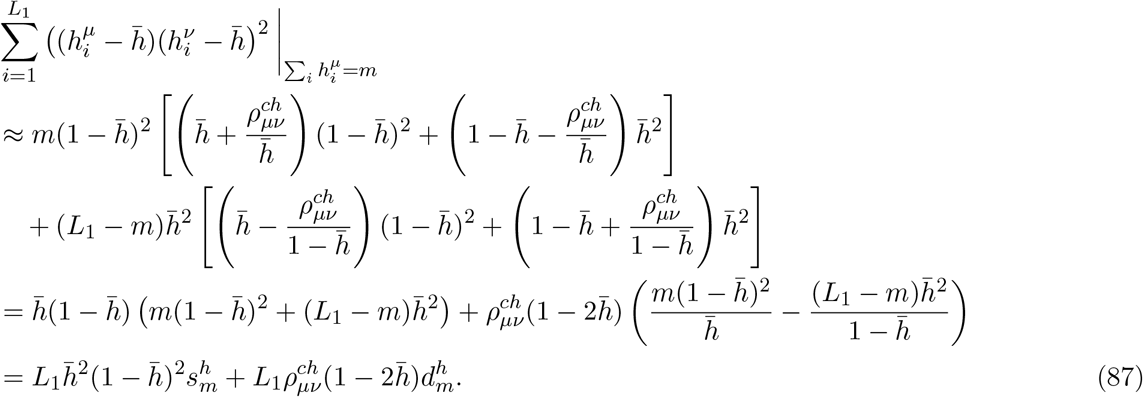

Here, we defined variable 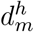 by

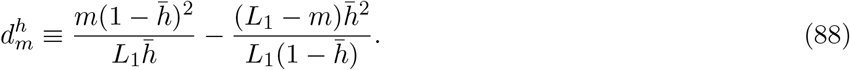

From a parallel argument, we have

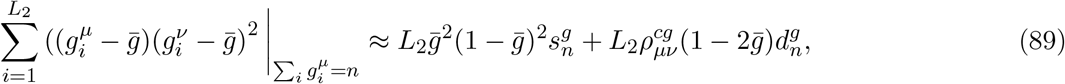

where

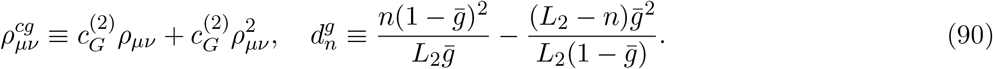

The second term of Eq. 85 is rewritten as

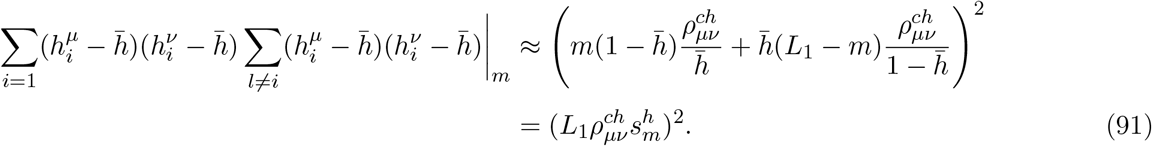

This means that the noise term is written as

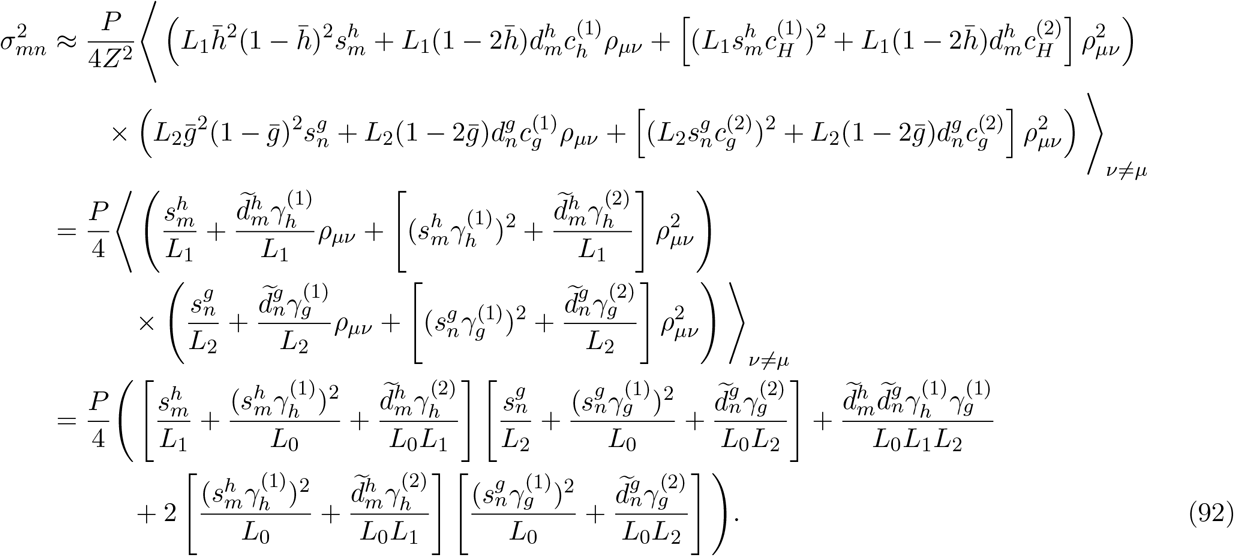

Here, for brevity, we introduced following variables:

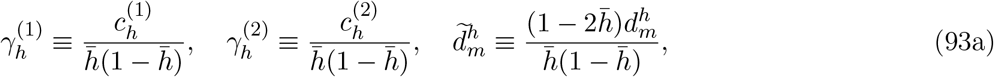

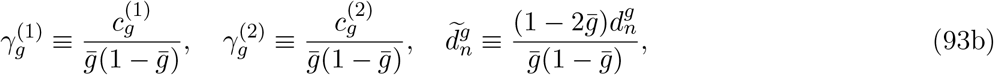

and used 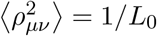 and 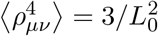.

### 4.4 Implementation Details

#### 4.4.1 Figure 1: Analytically tractable deep and shallow perceptron models

In Figs. 1F–J, we estimated capacity by evaluating classification performance for *P* = [*P*_min_, 2*P*_min_, …, *P*_max_] under baseline parameters *b* ∈ [0.0, 0.05, 0.1, …, 2.0], and then identifying the largest number of patterns for which the model achieved 90% classification accuracy under the optimal baseline parameter. We evaluated classification accuracy by averaging over 10 random seeds. The choice of a 90% threshold is somewhat arbitrary, but the results were qualitatively consistent when using a 95% threshold. We did not use 100% accuracy as the threshold because the model often failed to achieve exact 100% accuracy due to finite-size effects. In the one-layer model with *L*_1_ intermediate-layer neurons, we set *P*_min_ = max{0.005*L*_1_, 1} and *P*_max_ = *L*_1_. In the two-layer gated model with *L*_1_ and *L*_2_ intermediate-layer neurons, we set *P*_min_ = 0.005*L*_1_*L*_2_ and *P*_max_ = 0.5*L*_1_*L*_2_. We set the input-layer size to *L*_0_ = 100 in all settings.

The orange analytical curves in panels E–H and the viridis curves in panel D were estimated using Eq. 68, whereas the blue curves in panels E–H were estimated using Eq. 75. In panel H, for the 2L-gated model with *L*_*y*_ output units, we set the shared first intermediate-layer width to *L*_1_ = 800*/*(*L*_*y*_ + 1), and used *L*_*y*_ second intermediate-layer modules, each containing *L*_2_ = 800*/*(*L*_*y*_ + 1) neurons dedicated to one output neuron. In the 1L-vanilla model, we set *L*_1_ = 800.

#### 4.4.2 Figure 2: Comparison with deep neural networks

For the two- and three-layer neural networks described in Methods 4.2.3, we followed a standard deep neural network training protocol. Specifically, we used two output neurons (corresponding to labels 0 and 1), and trained the networks using an Adam-optimizer-based implementation of backpropagation with cross-entropy loss. We set the mini-batch size to *P/*100 (where *P* is the number of patterns) and trained the networks for 1000 epochs. In Fig. 2C, we plotted learning curves for *η* = 10^−4^, 3 × 10^−4^, 10^−3^, while in Figs. 2B and D we used the optimal learning rate among these values. As before, capacity was estimated using a 90% accuracy threshold and the average accuracy over 10 random seeds.

#### 4.4.3 Figure 3: Omniglot character classification task

Using the 128-dimensional image representations obtained from the pre-trained network (detailed in Methods 4.1.2), we trained the two-layer gated, one-layer vanilla, and two-layer vanilla networks on the 15 × 1024 training images for binary classification. Each of the 1024 characters was assigned a random binary label, and the network was trained to associate the training images of each character with its corresponding label.

For both the two-layer gated and one-layer vanilla networks, we simulated the models across a range of intermediate-layer activity sparsity values and selected the sparsity level that maximized generalization performance. More specifically, we selected the sparsity parameter *ρ*_*sp*_ from {0.025, 0.05, …, 0.5}, and set the bias parameter of each hidden neuron as

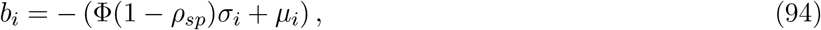

where *µ*_*i*_ and *σ*_*i*_ are the mean and standard deviation of the inputs to neuron *i* over the training data, and Φ(*x*) is the cumulative distribution function of the Gaussian distribution. In the two-layer gated network, we used the same sparsity parameter *ρ*_*sp*_ for both the first hidden-layer activity *h* and the gating input *g*. In addition, we sparsified the random fixed weights in the two-layer gated and one-layer vanilla networks using a connection probability of 0.3. This choice was motivated by our observation that sparse connectivity improves generalization (Fig. 1H), consistent with previous work [13].

In panels C and D, we set the intermediate-layer width to *L*_*h*_ = 1000 for the two-layer gated and two-layer vanilla models, and to *L*_*h*_ = 2000 for the one-layer vanilla model. We measured the performance shown in panel E after 10 epochs of training. In panel F, where images were presented directly to the networks, we rescaled the input images to 35 × 35.

#### 4.4.4 Figure 4: RL tasks

In both the Reacher and Pendulum tasks, we used a two-layer feedforward critic network with `tanh` activation, while the actor network was implemented as either a two-layer gated, one-layer vanilla, or two-layer vanilla network. For all three actor architectures, we used the `tanh` activation function instead of the step function. We set the number of neurons in each hidden layer to *L*_*h*_ = 64, except for the one-layer vanilla actor network, for which we used *L*_*h*_ = 128 so that the total number of hidden neurons was matched across architectures.

We used the PPO algorithm [45] with generalized advantage estimation [71], a standard approach for deep reinforcement learning in continuous action spaces [46]. After each trial, we updated the parameters of both the critic and actor networks. For the critic network and the two-layer vanilla actor network, all weights were updated by backpropagating the error. By contrast, in the two-layer gated actor network, we updated only the weights between the two hidden layers. Because the projections from the second hidden layer to the output were fixed, this is equivalent to directly projecting the error signal onto the connections between the two hidden layers. Similarly, in the one-layer vanilla network, we updated only the weights between the hidden layer and the output layer.

We simulated each network over a range of learning rates while keeping the other hyperparameters fixed, and then selected, for each architecture separately, the learning rate that maximized cumulative reward. Please see the code for further details.

#### 4.4.5 Figure 5: Cerebellar circuit model

In the cerebellar circuit model,we set the number of mossy fiber (MF) axons to *L*_*MF*_ = 100, granule cells (GC) to *L*_*GC*_ = 400, Golgi cells to *L*_*gol*_ = 50, Lugaro cells (LG) to *L*_*LG*_ = 200, stellate cells (SC) to *L*_*SC*_ = 200, Purkinje cell (PKC) branches to *N*_*branch*_ = 50, Purkinje cells to *L*_*PKC*_ = 8, basket cells (BC) to *L*_*BC*_ = 200, and cerebellar nuclei (NC) neurons to *L*_*NC*_ = 1 unless otherwise specified. Except for panel G, the number of synaptic inputs from mossy fibers to each granule cell was set to *K*_*GM*_ = 5 in the E/I model and *K*_*GM*_ = 10 in the model without E/I constraints. Similarly, we set the number of granule-cell inputs to each stellate cell to *K*_*SG*_ = 10, and the number of stellate-cell inputs to each Purkinje cell dendrite to *K*_*PDS*_ = 5.

Except for panel H, we set the bias parameter of stellate cells using a target sparsity of 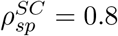. Specifically, the bias parameter of the *i*-th stellate cell was set as

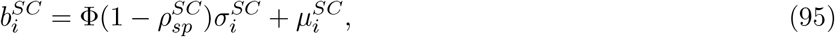

where 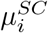 and 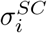 are the mean and standard deviation of the inputs to the *i*-th stellate cell. As in Figures 1 and 2, we optimized the baseline of granule cell activity and the threshold for shunting inhibition at each Purkinje cell dendrite to maximize capacity and defined the capacity as the maximum number of patterns for which the network achieves 90% classification accuracy.

In the ablation model with adaptation depicted in panel E, we adjusted the baseline parameter *b* of target neurons receiving direct input from the ablated neurons. For example, in the case of Golgi cell ablation, we adjusted the granule cell baseline *b*. Note that this adaptation does not recover capacity, because the sparsity of mossy fiber inputs varies across input patterns, which biases the overall granule cell activity level in the absence of balancing inhibition from Golgi cells.

In panel I, we set the number of patterns to *P* = 10000 and used the baseline parameter *b* = 0.65 that maximizes the classification accuracy. Note that in this model setting, all Purkinje cells receive a teaching signal from the same inferior olive neuron (though it doesn’t need to be the case); hence, we picked random cell pairs for the right panel of Fig. 5I.

### 4.5 Code availability

Codes are made publicly available at https://github.com/nhiratani/deep_cerebellum. Generative AI tools (ChatGPT & Claude Code) were used for assistance in writing and coding, but the author takes full responsibility for all the contents.

## 4.6 Acknowledgements

This work was partially supported by the McDonnell Center for Systems Neuroscience.

